# Integrating Alternative Fragmentation Techniques into Standard LC-MS Workflows Using a Single Deep Learning Model Enhances Proteome Coverage

**DOI:** 10.1101/2025.05.28.656555

**Authors:** Nikita Levin, Cemil Can Saylan, Joel Lapin, Yana Demyanenko, Kevin L. Yang, John Sidda, Alexey I. Nesvizhskii, Mathias Wilhelm, Shabaz Mohammed

## Abstract

We built and characterised a mass spectrometer capable of performing CID (both beam type and resonant type), UVPD, EID and ECD in an automated fashion during an LCMS type experiment. We exploited this ability to generate large datasets through multienzyme deep proteomics experiments for characterisation of these activation techniques. As a further step, motivated by the complexity generated by these dissociation techniques, we developed a single Prosit deep learning model for fragment ion intensity prediction covering all of these techniques. The generated model has been made publicly available and has been utilised in FragPipe within its MSBooster module. Rescoring allowed both data-dependent acquisition (DDA) and data-independent acquisition (DIA) to achieve on average more than 10% increase in protein identifications across all dissociation techniques and enzymatic digestions. We demonstrate that these alternative fragmentation approaches can now be used within standard data analysis pipelines and can produce data competitive to CID in terms of efficiency, but in the cases of EID and UVPD with far richer and more comprehensive spectra.

## Introduction

Mass spectrometry is the key approach for proteomics analysis^1^ due to its specificity, sensitivity and speed. Information corresponding to the sequence is acquired by dissociating each proteolyzed protein peptide into fragments via an established process, recording the spectrum and identification by specialist software.^1^Collision induced dissociation (both beam type and resonance based) has endured as the gold standard for peptide sequencing on account of its fragmentation explainability, reproducibility and outstanding speed.^2^ A variety of alternative dissociation techniques have been proposed as fragmentation techniques complementary to CID. These include, but are not limited to, ultraviolet photodissociation (UVPD) using photons of various wavelengths^3–6^ and irradiation by electrons in a wide range of electron energies, such as electron capture dissociation (ECD),^7–10^ hot-ECD,^8,10,11^ electron transfer dissociation (ETD),^12–14^ and electron ionisation dissociation (EID).^15–17^ The electron-based methods, combined under an umbrella term ExD, differ from each other in the energy of electrons used and types of fragments generated. These techniques provide access to alternative fragmentation pathways that improve peptide sequence coverage and can allow for keeping CID-labile modifications intact. Among them, ETD and electron transfer/high-energy collisional dissociation (EThcD)^18^ were adopted in Tribrid Orbitrap instruments, advancing proteomics studies to a more accurate analysis of phosphorylated^19,20^ and glycosylated^21–25^ peptides. However, all of the alternatives (to CID) approaches have experienced anaemic uptake due to a catch-22 scenario: weak demand leading to less vendor attention on the development and subsequent delivery of complicated, compromised, low-efficiency and low-speed designs. As these techniques remain a niche, the motivation to build suitable data analysis software further exacerbate and lag against classical CID approaches.

In recent years, there has been significant progress in the development of more accurate tools for analysis of bottom-up proteomics data.^1^ In particular, it has been shown that using data-driven rescoring pipelines^26–30^ based on predictions of peptide properties such as their retention time or fragmentation spectra greatly increase numbers of identifications compared to standard database searches.^31^ However, while the focus of the proteomics community has been mainly set on developing experimental and analysis workflows for collisional data, the development of tools for UVPD and ExD data analysis is precluded by the scarcity of publicly available datasets for calibration of search algorithms. The generation of such data, in turn, requires a specialised, often custom, MS instrumentation. Recently, Papanastasiou and co-workers reported a novel instrument, Omnitrap, which represents a segmented ion trap allowing multiple ion-activation approaches within one mass spectrometry platform.^32^ In collaboration, we built an Exploris-Omnitrap hybrid instrument that allows access to both common types of CID and multiple laser– and electron-based dissociation techniques and demonstrated it for top down proteomics.^33^ Herein, we further developed this instrument to operate on an LCMS time scale in an automated fashion. We set out to generate a large dataset of UVPD, EID and ECD fragmentation of peptides which we will use to develop prediction models using the Prosit deep learning model^27^ with the intention of improving data analysis through spectra rescoring and opening up these dissociation techniques to newer MS operation approaches such as data-independent acquisition (DIA).

## Results

### Development of Omnitrap UVPD, ECD and EID LCMS methods

The results of our recent development and characterisation of UVPD, EID and ECD on the Omnitrap platform^33^ suggested that our configuration could be deployed in an LCMS data-dependent fashion for the analysis of complex peptide mixtures. As the conditions in direct-infusion experiments from our earlier work, such as number of available ions, injection times and ion transfer logistics, are typically more relaxed compared to automated LCMS analysis, an investigation is required to determine the optimal parameters for all dissociation techniques in LCMS experiments. Direct-infusion experiments reported previously^33^ were focused on higher resolution and signal/noise with no regard to duty cycle. As acquisition of spectra with this Orbitrap-Omnitrap configuration has limited parallelization potential (Supplementary Fig. S1a), we initially concentrated on reducing the scan length to increase the number of recorded spectra as it is necessary to handle the complexity of proteomes. The Omnitrap design requires ions to be cooled through a gas pulse prior to any ion manipulation. The original design used a single gas valve that had a maximum repetition rate of 10 Hz (Supplementary Fig. S1b). In order to improve the maximum rate of the Omnitrap we implemented the use of two valves, operating alternately, for gas injection, which results in potentially doubling speed (Supplementary Fig. S1b). Subsequently, we optimised the direct current (DC) potentials for ion transfer within the Omnitrap to reduce the background collisional fragmentation (Supplementary Notes, Supplementary Fig. S2, S3). We then focussed on increasing the numbers of identifications in LCMS experiments through application of pragmatic parameters for spectra acquisition (Fig 1a). We began with the characterisation of UVPD. As there are two parameters of UVPD that can be changed, namely, the number and energy of laser pulses, we first varied the number of laser pulses at a fixed energy of 3 mJ/pulse and then varied the energy for the same number of pulses. For data analysis, we started with using only *b* and *y* ions for identification, as they were previously shown to be the most abundant in UVPD of tryptic peptides.^6,34,35^ The analysis shows that increasing the number of laser pulses leads to greater numbers of identified PSMs and peptide sequences until a maximum is reached at four pulses (Fig. 1b). Further increasing the number of laser pulses used for dissociation results in an insignificant drop of identification rates, either due to secondary fragmentation or reduced scan rate. We selected four pulses for further investigation and varied the energy of each pulse. In this series of experiments, the maximum of identified PSMs and peptide sequences was observed at distinct energies depending on the type of fragment ions used for identification (Fig. 1c). Please see Supplementary Table 1 for structures and definitions of all fragment ions considered in this work. Using only *b* and *y* fragments, the maximum is observed at 5 mJ/pulse, while when other types of fragments characteristic of UVPD are employed, namely *a,c,x,z*,^4^ the maximum is located at 6 mJ/pulse. As we reasoned that *a,c,x,z* in contrast to *b*,*y* are more unique to UVPD, we opted for using 6 mJ/pulse in future experiments.

**Figure 1.**
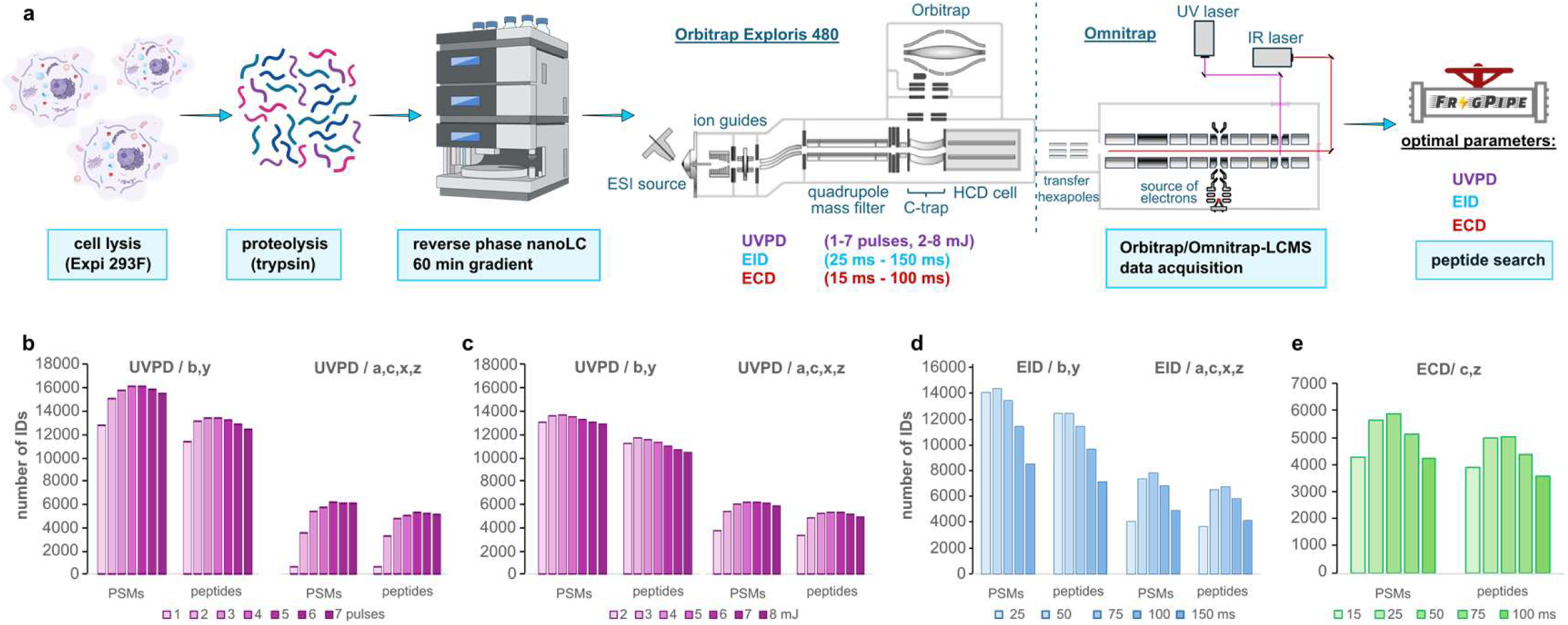
Optimization of ECD, EID, and UVPD parameters in bottom-up experiments. **a**, Experimental workflow; **b-e,** numbers of peptide-spectrum matches (PSMs) and peptides identified in UVPD experiments using different numbers of UV laser pulses at 3 mJ/pulse (b), UVPD experiments using four laser pulses at different energies of each pulse, (c) EID experiments using different irradiation times at 25 eV of electron energy (d), ECD experiments using different irradiation times at ∼1eV of electron energy (e). In UVPD and EID, *b*,*y* or *a,b,c,x,y,z* fragments were used for data analysis, *c* and *z* ions were using in the analysis of ECD data.

Next, we studied the optimal reaction times for EID. In typical ExD experiments, ions are transferred into the reaction chamber and undergo irradiation by electrons emitted by a heated filament^32^ during a specified amount of time (Supplementary notes, Supplementary Fig. S4). In EID experiments, we varied the irradiation times from 25 ms to 150 ms and measured the numbers of identified PSMs and peptides. We observed that *b* and *y* ions can be the most prominent in EID. When using just these two ions for analysis, the numbers of PSMs and peptides reach their maximum values at 50 ms of irradiation (Fig. 1d). At longer irradiation times, these numbers start to drop. Interestingly, the profile of peptide identification shows a much more distinctive dependence on the type of ions used for analysis compared to UVPD (Fig. 1d). At shorter irradiation times, *a,c,x,z* fragments are underrepresented compared to those of *b,y*, and the largest number of PSMs and peptides was observed at 75 ms (Fig. 1d). For the sake of absolute numbers of identifications, we chose to continue with 50 ms irradiation time. Finally, we optimised ECD where we used *c* and *z* type fragments for the data analysis and found 50 ms of irradiation to be optimal (Fig. 1e). We did not investigate other main-series types of fragments, as the absolute majority of products produced in ECD of relatively short peptides are *c* and *z* type ions.^7,8^ Since ECD is known to be a charge-dependent process favouring higher charge states, the value of 50 ms obtained using mainly doubly charged and less frequently triply charged precursors of tryptic peptides can be considered rather conservative. In order to characterize the fragmentation behaviour of ECD, UVPD and EID, a larger and more diverse range of peptides is required.

### Large-scale multi-enzyme Omnitrap-LCMS analysis

We increased diversity through use of more proteases, and we increased peptide depth by utilising offline fractionation (Fig. 2a). We chose trypsin, LysC, GluC, chymotrypsin and LysN as they have been shown to produce complementary results in terms of peptide length, protein sequence coverage, and frequencies and positions of amino acid residues across the peptide backbone.^36^ Next, we fractionated each digest^37^ into 20 pooled fractions and subjected all to ECD, EID, beam type CID (referred to as higher-energy collision induced dissociation or ‘HCD’ on Thermo instrumentation) and UVPD LCMS analyses. The choice of LC gradient time for the dissociation techniques was based on their maximal sequencing rate and to ensure they all produced similar numbers of scans.

**Figure 2.**
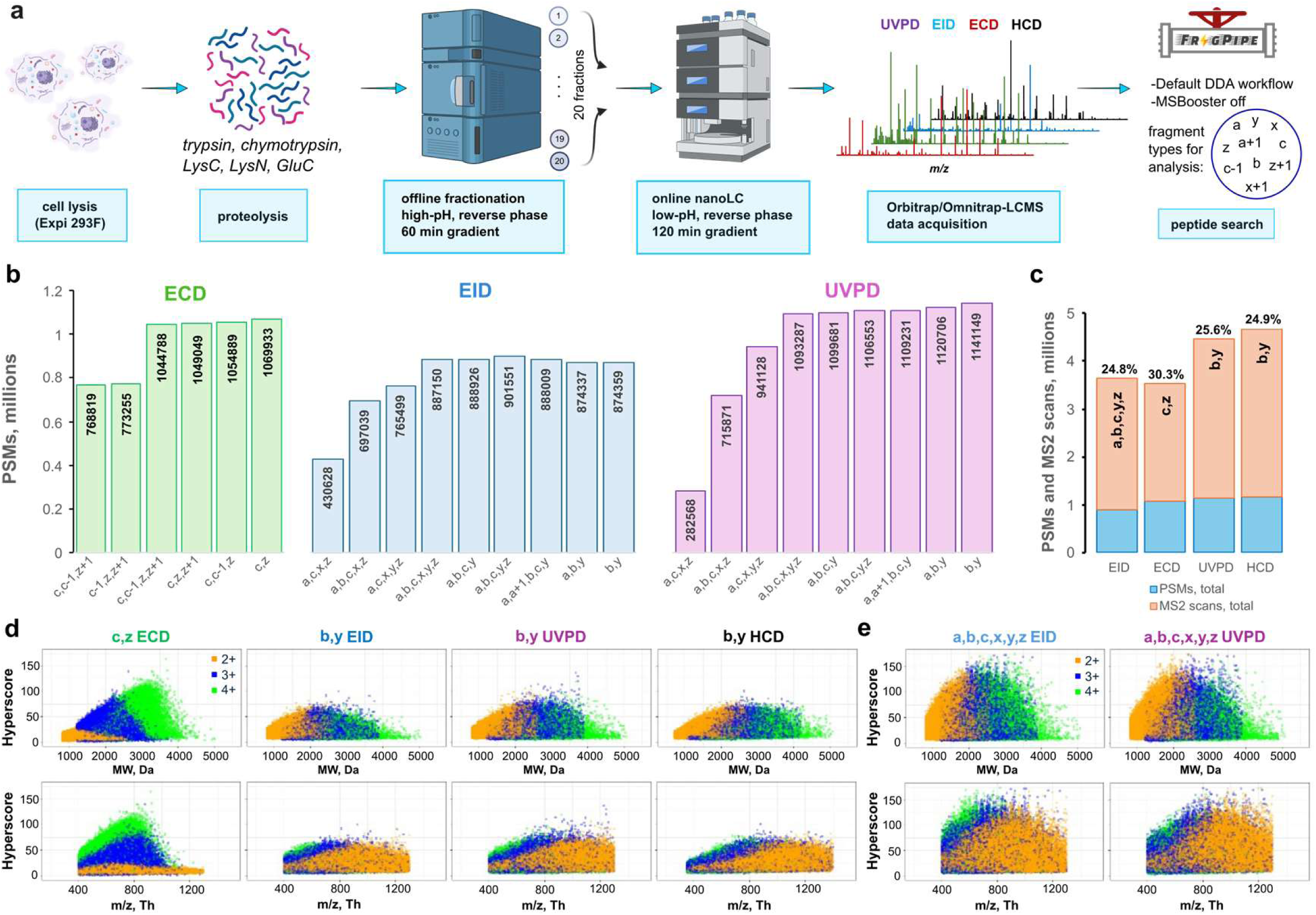
Large-scale bottom-up ECD, EID, UVPD, and HCD analysis. a) Experimental workflow; b) total numbers of peptide-spectrum matches (PSMs) in ECD, EID, and UVPD experiments identified using different combinations of fragment types; c) highest numbers of PSMs from b) (blue) and total numbers of acquired MS2 scans (orange) in ECD, EID, UVPD, and HCD experiments; the rate of PSM identification is shown above each corresponding bar; d) scatter plots of hyperscore distributions of 2+, 3+, and 4+ charge states of unique PSMs (unique combination of amino-acid sequence, charge, and modification selected by highest hyperscore) acquired in ECD using *c* and *z* fragments and in EID, UVPD, and HCD using *b* and *y* fragments; e) scatter plots of hyperscore distributions for each unique PSM acquired in EID and UVPD using *a,b,c,x,y,z* fragments.

The analysis of UVPD, EID and ECD data is not as straightforward as that of HCD data. The major products of HCD are well characterised, with *a*,*b*,*y* ions dominating data. In contrast, UVPD and EID are known to produce all main-series types of peptide fragments as well as some radical *a+1* and *x+1* ions,^4,15,38^ with the latter two largely understudied. The average proportions of each type of main-series fragments have been reported for UVPD;^6,34,35^ however, the effects of using these ions and their combinations in the automated data analysis have not been extensively discussed. We therefore analysed the acquired raw data using several unique combinations of the expected fragment types with the goal to maximise the numbers of identified PSMs while maintaining the same 1% false discovery rate (FDR). For ECD, the most important ions for robust identification were *c* and *z* (Fig. 2b). The addition of *c-1* or *z+1* had a minimal and slightly detrimental effect. Analogously, *b* and *y* were the dominant ion types for both EID and UVPD. However, *a, a+1, c, z* ions were beneficial for improving identification rates for EID, while *b, y* showed the best results in UVPD. The numbers when broken down to individual enzyme are similar to the global result, although tryptic and LysC peptides enhance the formation of *z+1* ions while impairing the formation of *c-1* in ECD and favour the generation of *y* ions in EID and UVPD compared to other enzymes (Supplementary Fig. S5). The results for the UVPD and EID seem to be strongly dependent on *y* ions and to a smaller degree on *b* ions. While no extensive literature exists for EID, our UVPD data agrees with previous findings. Others also found that *b* and *y* fragments are the most abundant types of ions in 193 nm UVPD of tryptic peptides, and the ion current of *y* fragments is approximately double of that of *b*.^6,34^ Similarly, *b* and *y* fragments dominate the spectra in 213 nm UVPD of tryptic peptides, and the average number of annotated *y* fragments is twice that of *b* ions.^35^

All four fragmentation techniques produced similar numbers of PSMs (Fig. 2c). In total, EID data had the least number of PSMs (approximately 900k), while UVPD, which has the fastest acquisition rate among all Omnitrap techniques studied here, had 1141k. Surprisingly, the charge-dependent ECD came closest to UVPD with 1070k PSMs, even though its scan rate was essentially the same as EID. HCD showed the highest numbers with 1160k PSMs. Pleasingly, the efficiency of peptide sequencing by EID and UVPD, expressed as the ratio between the numbers of PSMs and MS2 scans, is essentially the same as by HCD, and the efficiency of sequencing by ECD is higher by approximately 5% (Fig. 2c). This was surprising considering the relative inefficiency of ECD for doubly charged peptides which represent a substantial subset of identified peptides (Supplementary Fig. S6).

The MSFragger hyperscore (generated for each PSM) can serve as an indirect measure of the number of fragments found in a spectrum in relation to the number of fragments expected for that peptide, akin to a spectrum quality score.^39^ We plotted distributions of hyperscores of all unique precursors (*i.e.*, unique combinations of amino-acid sequences, charge states, and modifications, Supplementary Fig. S7) per charge state using *c,z* fragments in ECD and *b,y* fragments in UVPD, EID and HCD (Fig. 2d). Expectedly, distributions of hyperscores in ECD are strongly charge-dependent, with doubly charged precursor assigned with substantially lower values. Furthermore, the hyperscore distributions for 3+ and 4+ precursors in ECD have an apparent maximum at 800 Th. A similar trend was reported earlier by Good *et al.* for ETD of tryptic and LysC peptides, where the percent of bonds cleaved by ETD begins to drop at approximately 600 Th for 3+ precursors and 650 Th for 4+ ones.^13^ When analysing solely *b* and *y* ion series, EID, UVPD and HCD all show very similar hyperscore distributions for the same charge states of precursors (Fig. 2d). The upper boundaries of hyperscore distributions for these dissociation techniques start to drop beyond approximately 2500 Da. We interpret these observations as the reduction of the signal-to-noise (S/N) ratio that follows the spreading of available fragment signal across larger numbers of produced fragments in spectra of long and highly charged peptides, *i.e.* signal splitting. The difference in number of identifications with same 1% FDR was marginal for UVPD and EID when we increased the number of fragment types all the way up to *a,b,c,x,y,z*, as long as *b* and *y* fragments were included (Fig. 2b). We therefore investigated how choice of types of fragments for analysis affects hyperscores (Fig. 2e). Clearly, increasing the search space by adding more types of fragments results in greatly improved hyperscores for both EID and UVPD, indicating larger numbers of dissociated bonds and data rich spectra.

### Deep learning modelling of UVPD, EID and ECD fragment intensities

Following data collection, we asked the question whether we can employ advance data analysis techniques to improve search results by making use of the high values of hyperscores produced in ECD, EID and UVPD. Scoring can be remarkably boosted if peptide-spectrum matching is performed against experimental or *in-silico* generated spectral libraries.^31^ Deep learning models have demonstrated promising results in predicting CID-based spectra of peptides using just peptide sequence, charge state and collisional energy as input,^26–28,40^ but no such models exist for other fragmentation techniques due to lack of data for training. We therefore set out to use the datasets generated in this work to train a deep learning algorithm for prediction of EID, ECD, HCD and UVPD fragmentation spectra. Training a deep model requires turning the raw data into a dataset containing correctly annotated peak intensities. This implies that we need to solve potential clashes such as, for example, *a+1* ion, which is a radical *a* ion coupled with an additional hydrogen atom, *versus* the ^13^C peak for an *a* ion (Supplementary Tabe 1). For all datasets, we performed an automated annotation of major fragment types expected in EID, ECD and UVPD (Supplementary Table 1) in the Oktoberfest framework.^30^ The comparison of [*a+1*]/[*a*] ratio in HCD, EID and UVPD suggests that a high proportion of *a+1* in EID and UVPD spectra are ions originating from gas-phase electron– and photon-based chemistries (Fig. 3a, Supplementary Fig. S8-12 and Supplementary Notes). Annotated spectra in hand, we could define our model’s ion dictionary and curate training and validation datasets. The original Prosit model^27^ architecture was designed around a structured output space, *i.e. b* and *y* fragments with lengths 1-29 and charges +1, +2 and +3. By contrast, the model trained on our data has an unstructured output space, with fragment ions chosen based on frequency of occurrence (greater than or equal to 100 occurrences, Supplementary Fig. S9-12). Besides peptide sequence and precursor charge, the model also takes the categorical fragmentation type as input; since the HCD data was acquired on a single instrument, it was unnecessary to use collision energy as additional input to the model, as was performed for previous Prosit models.^27^ Our model shares the similarity with the original Prosit model insofar that the sequence and metadata are separately encoded into latent spaces and combined in the interior of the network, but the metadata has slightly changed, and the model outputs the intensities of 815 fragments of various length, charge and fragment type (Fig. 3b). Results show very little overtraining: the median Pearson correlations for ECD, UVPD, HCD and EID are 0.919, 0.931, 0.950 and 0.897, respectively, on the training set, and the corresponding scores for the test set are only ∼0.005 points behind for each fragmentation method (Fig. 3c, Supplementary Fig. S13, S14). We further observe that precursor charge is quite consequential for prediction performance, with precursor charges greater than 2 displaying an increasingly wide range of Pearson correlations, likely due to the sparsity of high charge precursors in the training set and increasingly complex fragment ions present in the spectra. Although the model outputs predicted intensities for fragments of all types, no matter the fragmentation method, we see that conditioned on the fragmentation method the model reliably assigns appreciable intensity only to those fragments expected for each fragmentation method, e.g. *b* and *y* for HCD and *c* and *z* for ECD (Fig. 3d,e). The model is also able to predict intensities of *b*, *y* and minor fragments, such as *a*, *a+1*, *x*, *x+1*, *c*, *z*, in UVPD and EID, although the predictions of low-intensity ions for the latter seem slightly less accurate (Fig. 3f,g). With this model in hand, these activation methods can now benefit from any state-of-the-art processing, be it data-driven rescoring, DIA or proteome-wide spectral library searching.

**Figure 3.**
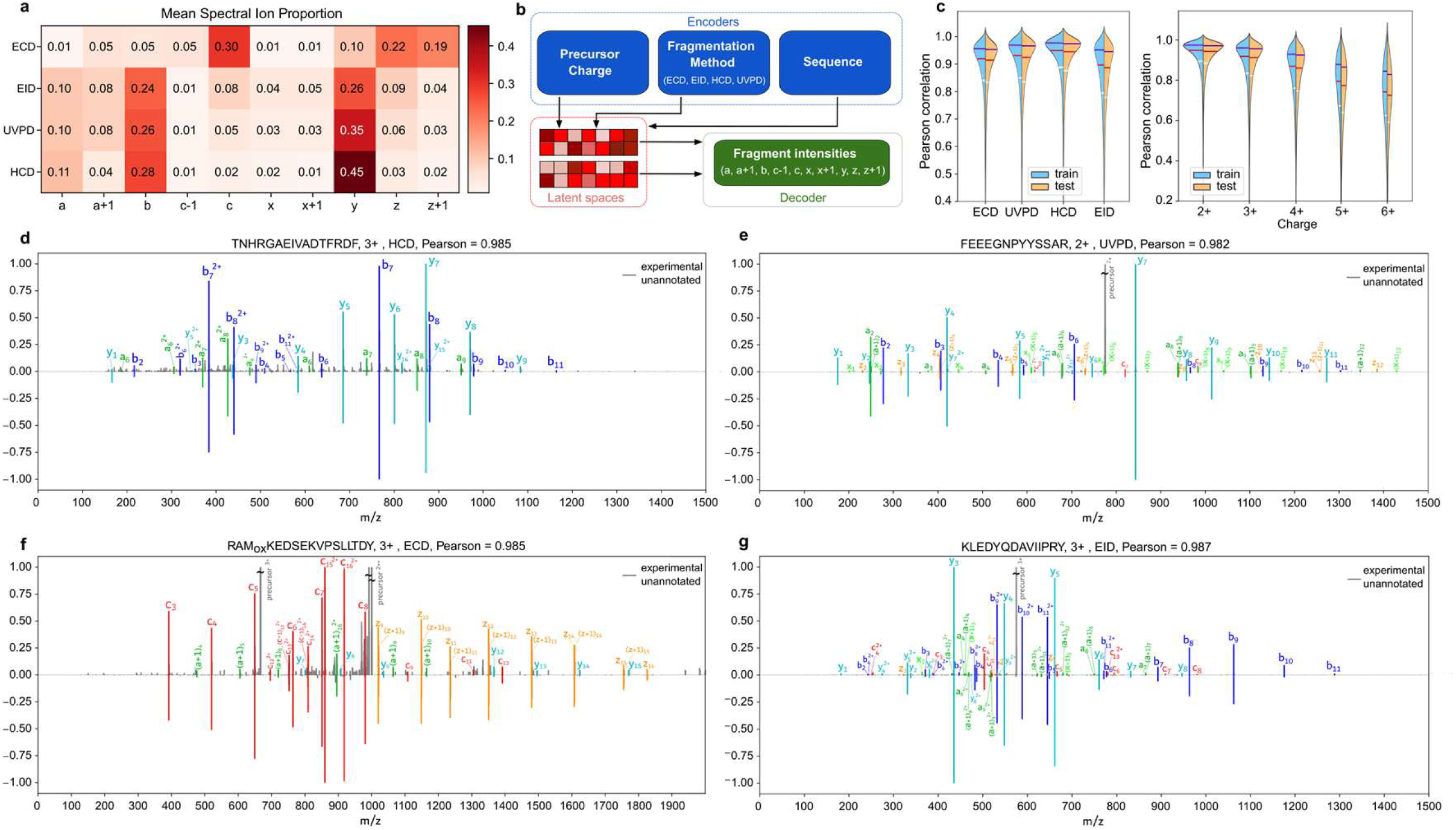
Deep learning training pipeline, from annotation to evaluation. **a**, Heatmap of mean fragment-ion proportions in ECD, EID, HCD, and UVPD spectra acquired across all enzymes. Annotation was performed for 10 ion types: *a*, *a+1*, *b*, *c-1*, *c*, *x*, *x+1*, *y*, *z*, *z+1* (Supplementary Table 1). **b,** The modified Prosit deep learning architecture for prediction of fragment ion intensities in ECD, EID, HCD, and UVPD spectra. The input parameters (peptide sequences, precursor charge state and fragmentation method) are encoded into a latent representation (latent space). This representation is then decoded to predict fragment ion intensities. **c,** Pearson correlation coefficients between predicted and experimental spectra in training (blue) and test (orange) sets separated by fragmentation method (left) and charge state (right). Horizontal white, red, and blue lines correspond to 25, 50, and 75% percentiles, respectively. Distributions protruding beyond 1.0 are plotting artefacts. **d-g,** Mirror plots of selected precursors in HCD (d), UVPD (e), ECD (f), and EID (g) data. Each mirror plot compares experimental (top) and predicted (bottom) fragment intensities, with each type of fragments uniquely colourised.

### Rescoring of the UVPD, ECD and EID data using fragment intensities predictions

An efficient control of FDR in database searching is critical for identification of true positive peptide matches. Previously, we showed that data-driven rescoring of CID data using the Prosit deep learning model greatly improved numbers and accuracy of peptide identifications.^27^ We hypothesized that predicting fragment intensities would be beneficial for improving the results of the database searches of UVPD, EID and ECD data as well. We started with a coarse and indirect estimate of the effect of predictions on the separation of target PSMs from decoys. Using the optimised MSFragger results we first calculated the ratios between the numbers of all observed and all possible theoretical fragment ions in each identified spectrum (Fig. 4a and Supplementary Fig. S15, upper distributions). The resulting distributions for target and decoy PSMs were heavily intermixed and shifted towards smaller ratios. EID and UVPD ratios were particularly small due to a large number of theoretical ions. We then calculated the same ratios but only allowed fragments predicted by Prosit (Fig. 4a and Supplementary Fig. S15, lower distributions). Including only predicted fragments split the distribution of ratios of target PSMs, where the majority shifted towards higher values with a larger portion being above 0.8, and the remainder were essentially unchanged. At the same time, the ratios of decoy PSMs remained clustered at lower values. This indicates a substantial improvement in alignment between the observed and the predicted fragment ions.

**Figure 4.**
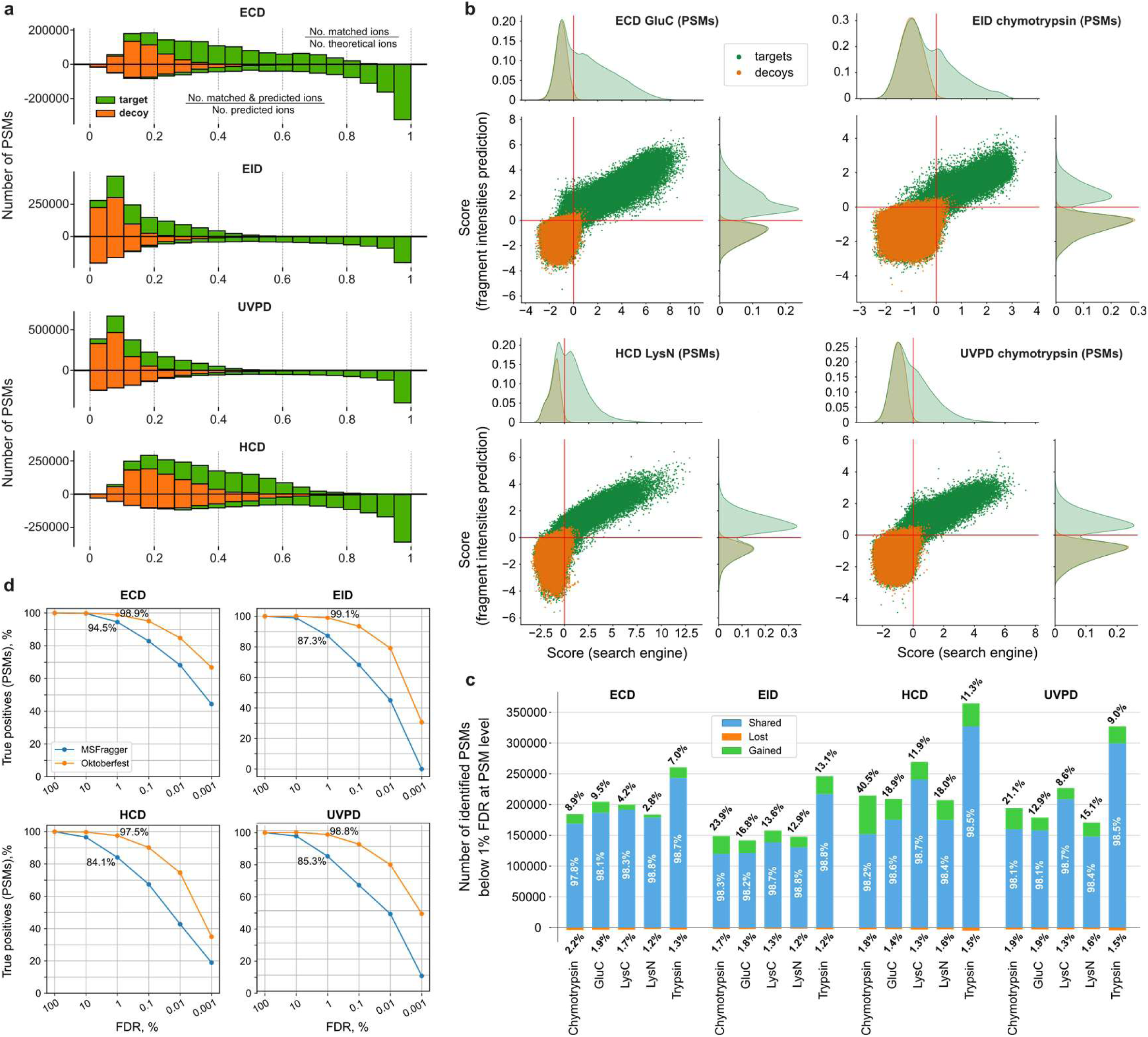
Intensity prediction improves database search quality of ECD, EID, HCD, and UVPD data. **a**, Histograms of the ratios of experimentally observed ions to all theoretically possible fragment (upper distributions); Histograms of the ratios of experimentally predicted and observed ions to all predicted ions (lower distributions). **b,** Correlation of Percolator scores for all target (green) and decoy (orange) PSMs obtained from the rescoring of the MSFragger (top) and Oktoberfest (right) sets of scores for selected combinations of enzyme and fragmentation technique. The red solid lines indicate the 1% PSM-level FDR cutoffs. For database searches scores, the best combinations of fragment types from Fig. 2b were used; for Oktoberfest scoring, most frequently annotated fragment types (>4% of all annotated ions across all spectra) were used for each dissociation method (Supplementary Fig. 8). **c,** Numbers of shared (blue), gained (green), and lost (orange) PSMs identified at 1% PSM-level FDR using the Oktoberfest set of scores compared to original MSFragger search for each fragmentation technique per enzyme. The numbers correspond to the data from **b** and Supplementary Fig. 16-19. **d,** Proportions of the numbers of true positive PSMs to the estimated maximum number of true positive PSMs acquired using original MSFragger and Oktoberfest scores at different values of PSM-level FDR for each fragmentation technique, all enzymes combined.

Next, we applied data-driven rescoring using the Oktoberfest framework, which benefits from the here-developed fragment intensity prediction model by generating fragment intensity-dependent scores rather than only relying on the presence or absence of fragments. In combination with Percolator,^41^ these scores are aggregated into a single score that maximizes the separation of correct and incorrect matches. The resulting Oktoberfest scores were then compared to the Percolator-derived scores from MSFragger (Fig. 4b and Supplementary Fig. S16-19), which do not include fragment intensity-based features. For MSFragger database searches, we chose the best combinations of ion types for each fragmentation method from Fig. 2b, and for rescoring in Oktoberfest we employed all the most frequently annotated types of fragments (>4% of annotated ions in a spectrum, averaged across all spectra) for each fragmentation technique (Supplementary Fig. S8). Both sets of scores were filtered to 1% FDR using Percolator.^41^ While rescoring led to remarkable separation of decoys from targets for the majority of enzyme-fragmentation method pairs (as exemplified in Fig 4b, Supplementary Fig. S16-19), ECD in general demonstrated sufficient separation in database searches, such that rescoring only delivers marginal improvements in identifications (Supplementary Fig. S16). This in part explains the highest identification rate identified for ECD in the initial database searches (Fig. 2c). We attribute this to the relative cleanliness of ECD spectra (Fig. 3a) that consist primarily of *c* and *z* type fragments, precursor ions and charge-reduced species, thus reducing chances for random false matches. Interestingly, ECD was the only technique where it was possible to discriminate the distributions of charge states among target PSMs, which reflects the distinct charge-dependent kinetics of this process (Supplementary Fig. S20). Using rescoring, we were able to salvage a substantial number of PSMs in all combinations of enzyme and dissociation method (quadrant II in Fig 4b and Supplementary Fig. S16-19). At the same time, a fair number of PSMs initially identified were discarded as false positives (quadrant IV in Fig 4b and Supplementary Fig. S16-19).

To evaluate how this separation of scores translates into gains and losses of PSMs and peptides, we evaluated the results of the database search and rescoring at both 1% PSM-level (Fig. 4c) and 1% peptide-level FDR (Supplementary Fig. S21). The number of gained PSMs varied depending on the enzyme and fragmentation method between approximately 3% and 40.5%, with chymotrypsin HCD data producing an exceptionally impressive gain of 40.5%. The latter observation is consistent with our previous findings.^27^ Remarkably, chymotrypsin was also the main beneficiary of rescoring in UVPD and EID data. This demonstrates the usefulness of rescoring for expanded search spaces characterised by increased number of possible charge states, allowed missed cleavages and reduced enzyme specificity, all of which are typical for chymotrypsin (Supplementary Fig. S6). Consistent with the score distributions (Fig. 4b and Supplementary Fig. S16-19), ECD enjoyed the least numbers of gained PSMs and peptides regardless of protease among all fragmentation techniques (Fig. 4c and Supplementary Fig. S21). To explore the reasons of the varying number of gains observed, we investigated the recovery of true positive (TP) PSMs at various FDR cutoffs. For that we compared the numbers of TPs identified in each method with the total possible number of TPs across a range of FDR thresholds before and after rescoring (Fig. 4d, Supplementary Fig. S22). At 1% PSM-level FDR, rescored ECD, EID and UVPD recovered more than 97% of possible TPs, while the original database searches extracted approximately 95% in ECD, 87% in EID, 85% in UVPD and 84% in HCD. At a stricter FDRs of 0.01%, the results after rescoring still captured over 75% of all estimated possible TPs, with ECD showing the highest proportion approaching 85%. At the same FDR level, initial base searches identified less than 70% of possible TPs in ECD and less than 55% in all other dissociation methods (Fig. 4d). The analysis shows that data-driven rescoring using the pan-activation Prosit model recovers essentially all TPs from the initial MSFragger search results.

The rescoring data provided an opportunity to inspect the efficacy of each enzyme and dissociation technique for proteome analysis (Supplementary Fig. S23a). Trypsin, as expected, identified the most PSMs, peptides and proteins for every fragmentation technique. Chymotrypsin demonstrated the next best result, with LysC and LysN a little further behind, replicating previous trends observed for CID and ETD data.^42,43^ The enzyme GluC clusters with LysN, appearing to be slightly superior or inferior depending on the dissociation technique. Average protein sequence coverage was similar for each fragmentation technique (Supplementary Fig S24). In order to assess complementarity at the protein sequence level we represented our data at the amino acid level. In general terms, when comparing the complementarity of trypsin against its alternatives, we saw substantial improvements in proteome coverage for all fragmentation techniques (Supplementary Fig S23b); in fact, the combined coverage for LysN, LysC, GluC and chymotrypsin was more than that for trypsin. It should be noted that a more exhaustive 2D LCMS analysis of the trypsin data would not significantly increase proteome coverage, and so the amount of analysis time between the other enzymes *versus* trypsin is not an important factor in the comparison. Further analysis of unique coverage for each fragmentation technique showed that UVPD produced the most amount of unique data, with HCD and ECD close behind, and EID the least (Supplementary Fig S24. However, UVPD had significant overlap with EID which might be a reason for weak unique proteome coverage result for EID (Supplementary Figure S23c).

### Application of DIA with all activation techniques and Pan-activation Prosit Model

The spectral prediction model created in this work is portable and freely available as instance “Prosit_2025_intensity_MultiFrag” at the Koina model repository,^44^ and can be interfaced within any software suite. We implemented our model within FragPipe as part of MSBooster.^29^ Combined with the optimisation of search parameters in Fragpipe, we can now perform both DDA and DIA data analyses for all activation techniques. The ability to now utilise these activation techniques with DIA approaches led us to create DIA methodologies/routines for the Exploris-Omnitrap. The change in ion population, both in terms of ion density and distribution of charge states, required adjustments of the acquisition parameters for each dissociation technique both at the Exploris and Omnitrap level. We carried out LCMS analyses on a tryptic cell lysate digests from Human, Arabidopsis Thaliana and Escherichia coli cells. We introduced the latter 2 types of cells to assess the universality of the prosit model. To optimise duty cycle we chose to use the ‘normal isolation window’ approach with MS1 range bound to retention time.^45^ MSBooster, using the Prosit Multrifrag model, increased identification rates at PSM-, peptide– and protein-level for all three cell types. Arabidopsis and the Human lysate samples enjoyed the largest improvements, trading top position depending on exact context. On average ECD had the lowest gains across all samples with the worst result being 1.0%, 1.7% and 3.0% at the three levels for E.Coli while EID enjoyed the largest improvements across all three types of samples with the best result being 31.4%, 20.9% and 22.6% at the three levels for the Arabidopsis sample (Fig. 5).

**Figure 5.**
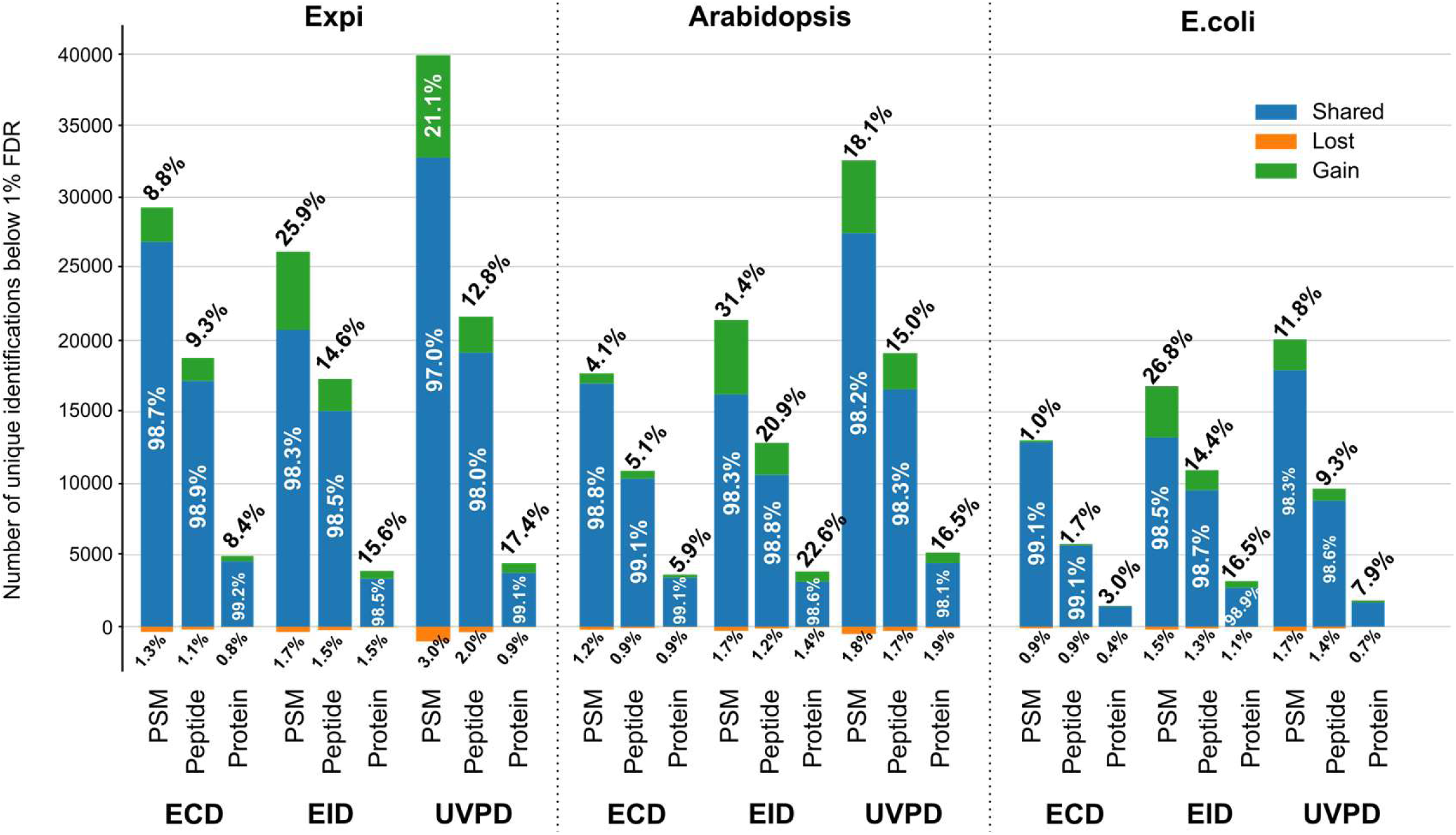
Intensity prediction improves search quality of ECD, EID and UVPD DIA data. Numbers of shared (blue), gained (green), and lost (orange) PSMs, peptides and proteins identified at 1% FDR with Prosit predictions implemented within MSBooster module of FragPipe in the UVPD, EID and ECD DIA data of unfractionated tryptic digests of Expi293F, Arabidopsis thaliana and E.coli proteins.

## Discussion

The intrinsic challenges of proteome characterisation have meant a necessary focus on sensitivity and speed to reach biologically useful levels of proteome information. In the last few years, the field has been able to reach a relevant level of proteome coverage in minutes and on sample sizes as low as a single cell.^46,47^ However, the great variety of protein/peptide physicochemical properties embedded within the proteome places a hard limit on what modern proteomics approaches based on trypsin and CID can potentially observe. The blind spots of these approaches are well described, and alternatives such as complementary proteases and activation techniques have been introduced although not widely adopted. In this work, we demonstrated that UVPD, EID and ECD can generate equivalent levels of data as HCD from the same numbers of spectra, *i.e.* these approaches can reach similar levels of efficiency on a proteome scale across the most popular enzymes. These data allowed us to train a model that can predict spectra approximately as accurately as those for CID. The advent of predicting spectra will now allow the community to consider if alternative dissociation techniques are more appropriate for their protein characterisation needs. An advantage of EID and UVPD is that their spectra are richer and less affected by the properties of the sequence intended to be studied and will certainly make more challenging proteoforms more accessible for characterisation. We demonstrated that our model could be used for the improvement of database search results both from DDA and DIA sources, creating software parity with cutting edge modern proteomics pipelines. The Exploris-Omnitrap configuration as described in this manuscript is a potent characterisation instrument that is arguably the most capable for the widest gamut of protein types, but it is not optimised for speed or sensitivity at speed. Our work shows the characteristics of each fragmentation method, and others^4,9,48^ have shown benefits. We hope that increasing the ability of the community to observe benefits of these activation approaches will increase demand and encourage vendors to consider implementation of more efficient designs allowing most if not all application types to be as successful on these activation platforms as the standard CID experiment, with the added advantage of far richer data and more confident characterisation.

## Methods

### Chemicals, solvents, cell lines and proteases

Sodium dodecyl sulfate (SDS), dimethyl sulfoxide (DMSO), chloroacetamide (CAA), acetonitrile, ethanol, triethylammonium bicarbonate (TEAB), ethyl acetate, formic acid and urea were purchased from Sigma Aldrich. Tris(2-carboxyethyl)phosphine (TCEP), ultrapure water and Pierce BCA protein assay kit were obtained from Thermo Fisher Scientific. Magnetic carboxyl coated magnetic beads were acquired from Cytiva. Proteases were purchased from: Wako (LysC), ImmunoPrecise Antibodies (LysN), Promega (trypsin and GluC), Roche (chymotrypsin). Expi293F (human cell line based on HEK 293) cells were purchased from Thermo Fisher.

### Cell lysis

Expi293F or E.coli cell pellets were resuspended in 2% SDS, 50 mM TEAB (for proteolysis using trypsin, LysN and LysC) or 1% SDS, 50 mM TEAB (for proteolysis using chymotrypsin) or 6 M urea, 50 mM TEAB (for proteolysis using GluC) and sonicated using BioruptorPico (Diagenode) for 30 cycles (30 s on/of). Protein concentration was determined using BCA assay following the manufacturer’s protocol. Proteins were then reduced and alkylated with 10 mM TCEP and 50 mM CAA for 30 minutes.

### Protein extraction

Ground *Arabidopsis thaliana* seedlings were homogenised in 100 mM Tris buffer pH 7.6 containing 5.6% SDS, 1x Protease inhibitor cocktail (Roche), 1x PhosStop (Roche), 10 mM TCEP, 20 mg/ml PVPP beads (Alfa Aesar) in an ice-cold ultrasonic water bath for 30 min. The homogenised extract was then incubated for 2h in an orbital shaker (400 rpm) at room temperature and clarified by two 10-min room-temperature centrifugation steps at 17,000 g.

### Proteolysis

In proteolysis using trypsin, LysC and LysN, proteins were digested using a modified single-pot solid-phase assisted sample preparation method (SP3) using magnetic carboxyl coated magnetic beads at 4:1 w/w bead-to-protein ratio.^49,50^ Proteins were precipitated onto the beads using acetonitrile (80% final concentration) and incubated for 30 min with shaking. The beads were washed three times with 80% ethanol, followed by three washes with 100% acetonitrile before incubation with digestion buffer (50 mM TEAB), containing a protease (trypsin, LysN or LysC) at 1:25 enzyme-to-protein ratio for 4 hours. Supernatant was collected, beads were washed with 2% DMSO solution, and the wash was combined with supernatant before acidification with formic acid to 5% final concentration and centrifugation at 16,000 x g for 10 minutes. In proteolysis using chymotrypsin, proteins samples were diluted two-fold and digested with chymotrypsin at an enzyme-to-protein ratio of 1:50 for 4 hours. Peptides were then acidified with formic acid and washed with saturated ethyl acetate five times.^51^ In proteolysis using GluC, protein samples were diluted to adjust urea concentration to 0.8 M and digested with chymotrypsin at an enzyme-to-protein ratio of 1:35 for 4 hours. As a final step in all protocols, peptides were desalted on Oasis HLB cartridges (Waters) and eluted with 50% acetonitrile in ultrapure water. The eluent containing peptides was dried using Genevac EZ-2.

### UPLC fractionation^37^

Digests of Expi293F proteins were reconstituted in water containing 5% formic acid and 5% DMSO and subjected to UPLC fractionation in a Water Acquity Plus system. The peptides were separated using a Waters ACQUITY UPLC peptide CSH C18 (1mm × 150 mm, 1.7 µm, 130Å) column. The composition of solvent B (80% acetonitrile, 20% water) changed from 12% to 40% over 55 minutes and then from 40% to 50% over 5 minutes. Solvent A comprised 10 mM TEAB (pH 8.0-8.2), 2% acetonitrile and 98% water. Each fraction was collected for 45 seconds (80 fractions in total). Fractions were concatenated as follows: fraction 1 was pooled with fractions 21, 41 and 61; other fractions were pooled in a similar fashion. Each pooled fraction contained approximately 6 µg of peptides in 200 µL of solvents. Each pooled fraction was then split into 10 aliquots, each aliquot was further diluted by adding 80 µL of 5% DMSO-5% formic acid to 20 µL of peptide mixture.

### LCMS analysis

LC-MS/MS data were acquired using an UltiMate 3000 nanoUHPLC system (Thermo Fisher Scientific) coupled either to an Orbitrap Exploris (Thermo Fisher Scientific) equipped with an Omnitrap (Fasmatech) or to an Orbitrap Ascend Tribrid (Thermo Fisher Scientific). The amount of material injected was 500 ng of each digest fraction except trypsin, for which 250 ng of each fraction were injected. For HCD analysis, the amount of material was further reduced by half for all proteases. The peptides were trapped on a C18 PepMap100 pre-column (300 µm i.d. x 5 mm, 100 Å, Thermo Fisher Scientific) using solvent A (0.1% formic acid in water), then separated on an in-house packed analytical column (50 µm i.d. x 50 cm in-house packed with ReproSil Gold 120 C18, 1.9 µm, Dr. Maisch GmbH). The composition of solvent B (0.1% formic acid in acetonitrile) changed from 10% to 33% over 30 min (for parameters optimisation), or from 10% to 33% over 60 min (for parameters optimisation and HCD analysis), from 8% to 28% over 120 min (UVPD, EID, ECD DDA; HCD DIA), or from 8% to 20% over 240 min (UVPD, EID, ECD DIA) at a flow rate of 100 nL/min. Full scan MS1 spectra were acquired in the Orbitrap (scan range 400-1300 m/z, resolution 60000, AGC target 300%). For DDA, top 20 (40 in HCD) most abundant peptides were selected each round for fragmentation. EID, UVPD and ECD DDA spectra were acquired in the Omnitrap. Their products were mass-analysed in the Orbitrap at 45000 resolving power, the AGC value was set to 200%, and the maximum injection time was set to 64 ms. ETciD DDA spectra were acquired in the linear ion trap of the Orbitrap Ascend. Products of ETciD were mass-analysed in the Orbitrap at 7500 resolving power, the AGC value was set to 200%, and the maximum injection time was set to 64 ms. HCD DDA spectra were acquired in the HCD cell of the Orbitrap Exploris. Products of HCD were mass-analysed in the Orbitrap at 7500 resolving power, the AGC value was set to 40%, and the maximum injection time was set to 64 ms. In DIA, scans were acquired using 4-or 8-m/z isolation windows in the 400-700 m/z range over 0-80 minutes, 500-700 m/z range over 80-160 minutes, and 600-900 m/z range over 160-240 min of a 240-min LC gradient. EID, UVPD and ECD DIA spectra were acquired in the Omnitrap. Their products were mass-analysed in the Orbitrap at 60000 resolving power, the AGC value was set to 2000%, and the maximum injection time was set to 50 ms. In DIA experiments, the amount of material injected was 400 ng of unfractionated tryptic digest of Expi293F cells, 1200 ng of tryptic digest of Arabidopsis thaliana and 300 ng of tryptic digest of E.coli.

### Database searches of DDA data

Spectra were searched using FragPipe (v22.0) MSFragger 4.1^39^ with standard ‘close’ search settings against Uniprot database (UPR_Homo sapiens_9606 accessed on 24.10.29). The number of missed cleavages was set to four when searching GluC and chymotrypsin data. Types of fragments searched by MSFragger were set in accordance with fragmentation techniques used in the corresponding experiments. Ions *a+1*, *c-1*, *x+1* and *z+1* were manually defined for searches. MSBooster^29^ and all quantification parameters were disabled in all searches including HCD data. Mass calibration and parameter optimizations were enabled. Data filtering was performed using Percolator (v3.6.5) and Philosopher (v5.1.1), followed by ProteinProphet. Results were reported using sequential FDR estimation, with a 1% FDR threshold at the protein level.

### Spectra annotation

Every raw file and its corresponding pepxml file generated by MSFragger were read and merged into a single dataframe, containing amino-acid sequence, precursor charge, modifications (if any) and raw mass spectrum. For each PSM identified by MSFragger, its corresponding raw spectrum was annotated by matching with 10 ppm tolerance against a theoretical spectrum containing all ions with the following parameters: ion types *a*, *a+1*, *b*, *c-1*, *c*, *x*, *x+1*, *y*, *z*, *z+1*, fragment charges +1 to +3, and fragment lengths 1-29. See Supplementary Table 1 for ion type nomenclature. The annotation was carried out excluding all matches in a 2.4 Da window centred on the precursor ion’s *m/z*. For additional quality control, we explicitly annotate the first four isotopic precursor peaks, for both the precursor charge and charge+1, for the purpose of excluding them from a coincidental identification with another ion. Annotations for all PSMs are saved to one dataframe per fragmentation method, along with overall statistics enumerating the occurrences of each searched ion from our permutation.

### Creation of deep learning datasets

Datasets for the training of models were created as multiple parquet files from the annotated PSM datasets for all fragmentation methods. The PSM list was first cleaned of decoy sequences and sequences that contain no identified annotations. The output space/ion dictionary of the model was determined by only including annotated ions that occurred 100 or more times in the dataset. The PSMs were de-duplicated to unique sequence/charge/fragmentation method instances, retaining the instance with the highest hyperscore from MSFragger. The resulting unique PSM dataset was then shuffled, split into train/validation/test and written to parquet files. For the purposes of multithreading during dataloading, the train set was written to 8-12 separate files, all approximately having the same number of samples.

All dataset parquet files contain the original raw file name, scan number, modified peptide sequence, charge, fragmentation method, annotated ion strings and annotated ion intensities. The raw file names and scan numbers are sufficient to map any training/evaluation example back to its raw spectrum. Peptide sequences, charges and fragmentation methods were the inputs to the model; the target outputs for training were constructed from the ion strings and intensities. With a defined ion dictionary for the model, each ion string was mapped to its designated index, and its intensity was placed into a vector with the same length as the number of ions the model predicts (815). For ions that were the same length or longer than the peptide, or whose charge was greater than the precursor charge, a “-1” was assigned as the intensity and used to mask out these ions for training. All other unannotated ions got an intensity of 0. All non-negative entries in the target spectrum vector were max-scaled by the most intense ion, such that the intensities fall between 0 and 1.

### Deep learning training and models

We used the original GRU cell RNN network of Prosit, but with specific architectural tweaks in order to accommodate the different metadata and the unordered structure of its output space. The metadata, *i.e.* charge and fragmentation method were both represented as one-hot vectors, concatenated and then linearly projected into 256 units; collision energy was not used as input for this model. Whereas the original Prosit model projected the metadata and sequence representation together multiplicatively, followed by replicating the combined representation into 174 identical vectors (Prosit’s original output length), our sequence dimension retained the original maximum sequence length dimension (30 amino acids) and was multiplicatively combined with the 256-long metadata vector. This encoding was then passed to the RNN GRU cell bi-directional decoder layer. The final regressor then applied a linear transformation to our 815-ion output space, followed by a LeakyReLU activation function and mean pooling over the sequence dimension, yielding an 815-long vector of intensity predictions.

The model was trained by masked spectral distance. All indices of ions impossible to occur for a particular peptide were multiplied by 0 so that they do not contribute to the gradient during training. All models were trained for 30 epochs. The Adam optimizer was used with a learning rate of 1e-4, linearly warmed up for 20,000 steps at the beginning of training. Pytorch was used for implementation. The model is publicly available via the Koina prediction service under the title “Prosit_2025_intensity_MultiFrag”.

### Rescoring with Oktoberfest

To assess rescoring under 1% FDR control at both the PSM and peptide levels, spectra were searched with varied parameters, utilizing the same versions of FragPipe, MSFragger and the fasta file as in the initial data analysis. The number of missed cleavages was maintained as previously established, with the following ion types specified according to the fragmentation method: *c*,*z* for ECD, *a*,*b*,*c*,*x*,*y*,*z* for EID, and *b*,*y* for both HCD and UVPD. To ensure compatibility and reduce confounding factors in the comparison, we modified several parameters: mass calibration, parameter optimizations, N-terminal acetylation, MSbooster, ProteinProphet and all quantification settings were disabled, and the maximum peptide length was set to 30. The “pin” files generated by FragPipe were concatenated and utilized for Percolator to obtain 1% FDR control at both the PSM and peptide levels. The results were subsequently used for downstream comparisons with the rescoring results.

Rescoring was conducted using Oktoberfest and Percolator (v3.6.5) on 100% FDR search results. No calibration for collision energy was applied, and iRT predictions were made utilizing the “Prosit_2019_irt” model. Intensity predictions were performed for the following ion fragmentations: *a+1*,*b*,*c-1*,*c*,*y*,*z*,*z+1* for ECD; *a*,*a+1*,*b*,*c*,*y*,*x*,*x+1*,*z*,*z+1* for EID; *a*,*b*,*y* for HCD; and *a*,*a+1*,*b*,*c*,*y*,*z* for UVPD. Percolator was used to filter the data to 1% FDR at both the PSM and peptide levels.

### Database searches of DIA data

Spectra were searched using FragPipe (v23.0) with MSFragger-DIA 4.1^52^ with standard ‘DIA_SpecLib_Quant’ search settings against Uniprot databases (“UPR_Homo sapiens_9606” accessed on 22.08.22, “UPR_ArabidopsisThaliana_3702” accessed on 22.08.22 and “UPR_EscherichiaColi” accessed on 22.09.07). EasyPQP library generation and DIA-NN quantification were turned of. Types of fragments searched by MSFragger were set in accordance with fragmentation techniques used in the corresponding experiments. Mass calibration and methionine oxidation variable modifications were turned on, and the maximum length of peptide was set to 30. The “Prosit_2019_irt” and “Prosit_2025_intensity_MultiFrag” models were used in MSBooster^29^ via Koina^44^ to predict peptide properties. Data filtering was performed using Percolator (v3.7.1) and Philosopher (v5.1.1), and results were reported at 1% PSM, 1% Peptide and 1% Proteins levels.

## Acknowledgements

N.L, Y.D and S.M. was supported by the EPSRC (V011359/1 (P)). J.S and S.M. was supported by BBSRC responsive mode (BB/T016272/1). A.I.N and K.L.Y were supported in part by NIH grants R01-GM-094231 and U24-CA210967 (to A.I.N.). J.L and M. W. was, in part, supported by an ERC Starting Grant (Grant No. 101077037). C.C.S was supported by the Elitenetzwerk Bayern (grant number F-6-M5613.6.K-NW-2021-411/1/1).

## Author contributions

N.L. and C. C. S. contributed equally. S.M. and M.W. conceived and supervised the project. N.L. and S.M. designed LCMS experiments and wrote the original draft. N.L. optimised parameters of Omnitrap-LCMS, performed LCMS experiments and initial data analysis. C.C.S. performed comprehensive data analysis and data post-processing. J.L. wrote the code for and inspected the Prosit Multifrag model. Y.D. and J.S. prepared samples. K.L.Y. and A.I.N. implemented the Prosit Multifrag model into FragPipe MSBooster. S.M., M.W. and A.I.N. acquired funding. All authors have revised the manuscript and have given approval to its final version.

## Supplementary notes

### Optimisation of DC gradients for ion transfer within the Omnitrap

For the characterisation of UVPD, we constructed an experiment using short 30 min LC gradients and used a tryptic cell lysate digest as the analyte. As there are two parameters of UVPD that can be changed, namely, the number and energy of laser pulses, we began with varying the number of laser pulses at a fixed energy of 6 mJ/pulse. In the analysis of the data acquired in UVPD experiments, we started with using only *b* and *y* ions for identification, as they were previously shown to be the most abundant in UVPD of tryptic peptides [55, 56, 57]. Please see Supplementary Table 1 for structures and definitions of all fragment ions considered in this work. The analysis shows that the numbers of identified peptide-spectrum matches (PSMs) and peptides reaches a maximum at two pulses and quickly starts to fall beyond that point (Supplementary Fig. 2a). As key characteristic of UVPD is the formation of *a,x,c,z* (in addition to *b* and *y*) ions, we asked the question if the trend was the same when considering these fragment types. The numbers of identifications indeed did follow the same trend but were remarkably lower than those obtained using *b* and *y* fragments (Supplementary Fig. 2b). Based on the earlier reports [55, 56, 57], we expected a higher proportion of identifications using *a,x,c,z* in our data. As *b* and *y* fragments can also be generated through ‘background’ collisional fragmentation which can occur in multiple locations during ion transfer, we investigated the origins of these ions. We repeated the experiment but now without triggering of the UV laser and observed a significant population of *b* and *y* ions. We found that the ion transfer to and within the Omnitrap was the source and so we tested several new approaches. In the original approach, ions would be transferred from the HCD cell of Exploris through the transfer hexapoles and into the Q2 segment of the Omnitrap (Supplementary Fig. 3a). Subsequently, the ions would be transferred into the Q8 segment for UVPD using an 8V DC gradient between Q2 and Q8. We tried two other designs of ion transfer for UVPD: a) use of a 4V gradient in the last ion transfer step and b) injection into the Q5 segments and then transfer to Q8 using a 4V DC gradient (Supplementary Fig. 3a). Comparing the numbers of PSMs, peptides, and proteins obtained we found that the third design allows reduction of the number of PSMs sequenced by the factor of 4 to negligible levels (Supplementary Fig. 3b). We therefore switched to this design and continued the general optimisation (see the article).

### Discussion and interpretation of annotation results

For all LCMS datasets, we performed an automated annotation of major fragment types expected in EID, ECD, and UVPD (Supplementary Table 1) in the Oktoberfest platform. After gathering annotation statistics over the entire dataset, we could establish an output ion fragment dictionary, based on the prevalence of any ion we searched for. This varies heavily between the fragmentation methods. For instance, while HCD is known to produce primarily *a*, *b* and *y* fragments, EID and UVPD, on average, contain appreciable proportions of *a*, *a+1*, *b*, *c*, *x*, *x+1*, *y*, *z*, and *z+1* (Fig. 3a). More detailed distributions of ion proportions per enzyme are given in Supplementary Fig. S8-12. Notably, the frequencies of these ions are very similar in EID and UVPD, with the former having on average more *c*, *x+1*, and *z*, and the latter having more *y* fragments. While *a+1* signal may in principle correspond to the ^13^C isotope of an *a* ion, the comparison of [*a+1*]/[*a*] ratio in HCD, EID, and UVPD suggests that a good proportion of *a+1* in EID and UVPD spectra are rather ions originating from gas-phase electron– and photon-based chemistries. The formation of *a+1* fragments in UVPD have been described in multiple publications [58], whereas in EID they have been largely ignored and to the best of our knowledge were only reported by Ly *et al*. [59] In ECD, radical *a+1* ions are believed to form *via* solvation of charge located at the backbone nitrogen (as opposed to solvation of charge at carbonyl required for the generation of *c*/*z* ions) [60]. In our data, they comprise on average 5% of all annotated signals which is almost the same proportion as in HCD. The frequency of *a* ions in ECD is however negligible (potentially random matches) as opposed to 11% in HCD (Fig. 3a), which confirms the non-vibrational nature of *a+1* in ECD. The high abundance of *z+1* ions in ECD can be explained by the presence of ^13^C isotopes of *z* fragments, but in this case we would expect the [*z+1*]/[*z*] ratio to be much lower at 35-50%, similar to [*a+1*]/[*a*] ratio in HCD. This suggests that *z+1* ions are possibly products of hydrogen atom transfer (HAT). The *c-1* fragments in ECD are far less frequent than *z+1* ions that they are believed to complement, the phenomenon also observed in ETD [61, 62]. Although its causes have not been discussed, a potential explanation can be given by reduced stability of N-centred radicals (assumed for *c-1* ions) [63], as compared to relatively stable C-centred radicals found in *a+1*, *x+1*, and *z* fragments [64]. Remarkably, *c-1* ions are not present in UVPD and EID data, and *z+1* fragments are very likely to be ^13^C isotopes of *z*, based on the [*z+1*]/[*z*] ratio (Fig. 3a, Supplementary Fig. S8). This suggests either non-ECD-like ion chemistry of their formation, or different energetics of the process reducing the propensity for HAT. Furthermore, *x* ions were previously reported to be by far the least abundant in UVPD of cationic peptides, hardly surpassing their frequency in HCD data (where they supposedly represent random matches) for different proteases [55]. Our data shows similar trend and suggests similar frequencies of *x* and *x+1*. EID contains the highest proportions of *x* and *x+1* fragments among all fragmentation methods, with the frequency of *x+1* even higher than that of *x* (Fig. 3a, Supplementary Fig. S8). Finally, HCD, EID, and UVPD are dominated by *b* and *y* fragments. Interestingly though, while the proportion of *b* ions is uniform across these three techniques and comprises on average approximately 25%, the average proportion of *y* ions tends to increase from the relatively modest 26% in EID through 35% in UVPD to nearly one half in HCD (Fig. 3a, Supplementary Fig. S8), potentially indicating different mechanisms involved. In ECD, *b* and *y* ions are relatively scarce and originate either from collisions with the buffer gas (*i.e.*, background CID) or, in the case of *y* ions, potentially from solvation of charge located at the backbone nitrogen as discussed above.

### Duty cycle of irradiation of precursor ions by free electrons in the Omnitrap

In typical Omnitrap ExD experiments, precursor ions are transferred into the reaction chamber and undergo irradiation by electrons emitted by a heated filament during a specified amount of time [65]. The efficient electron irradiation time, however, is only half of the time that the ions spend in the reaction chamber. This is due to the rectangular waveform potentials applied to the electrodes of the Omnitrap that alternate with radiofrequency (RF) [65]. These potentials allow electrons to enter the ion trap only during the positive RF phase and repel otherwise. We provide total ion confinement times for ExD throughout this paper, and the efficient irradiation times can be obtained by dividing these values by two.

---

**Supplementary Figure S1.**
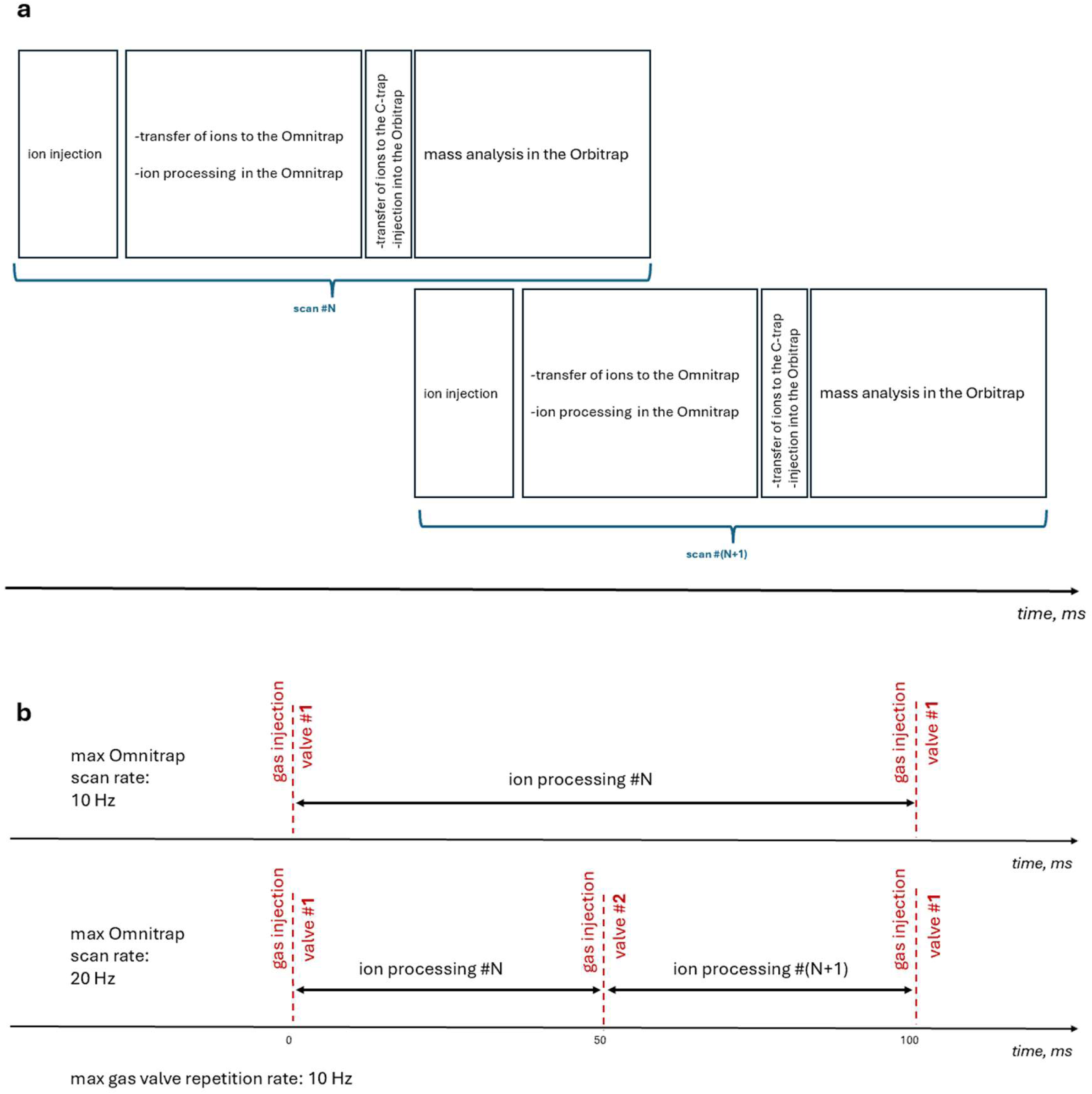
Ion processing in the Orbitrap-Omnitrap instrument. **a**, schematic workflow of ion processing in the Orbitrap-Omnitrap instrument; **b,** illustration of the minimum time distance between two injections of buffer gas into the Omnitrap using one valve (top) or two valves (bottom). The buffer gas is necessary for transfer, thermalisation, and trapping of ions.

**Supplementary Table 1.**
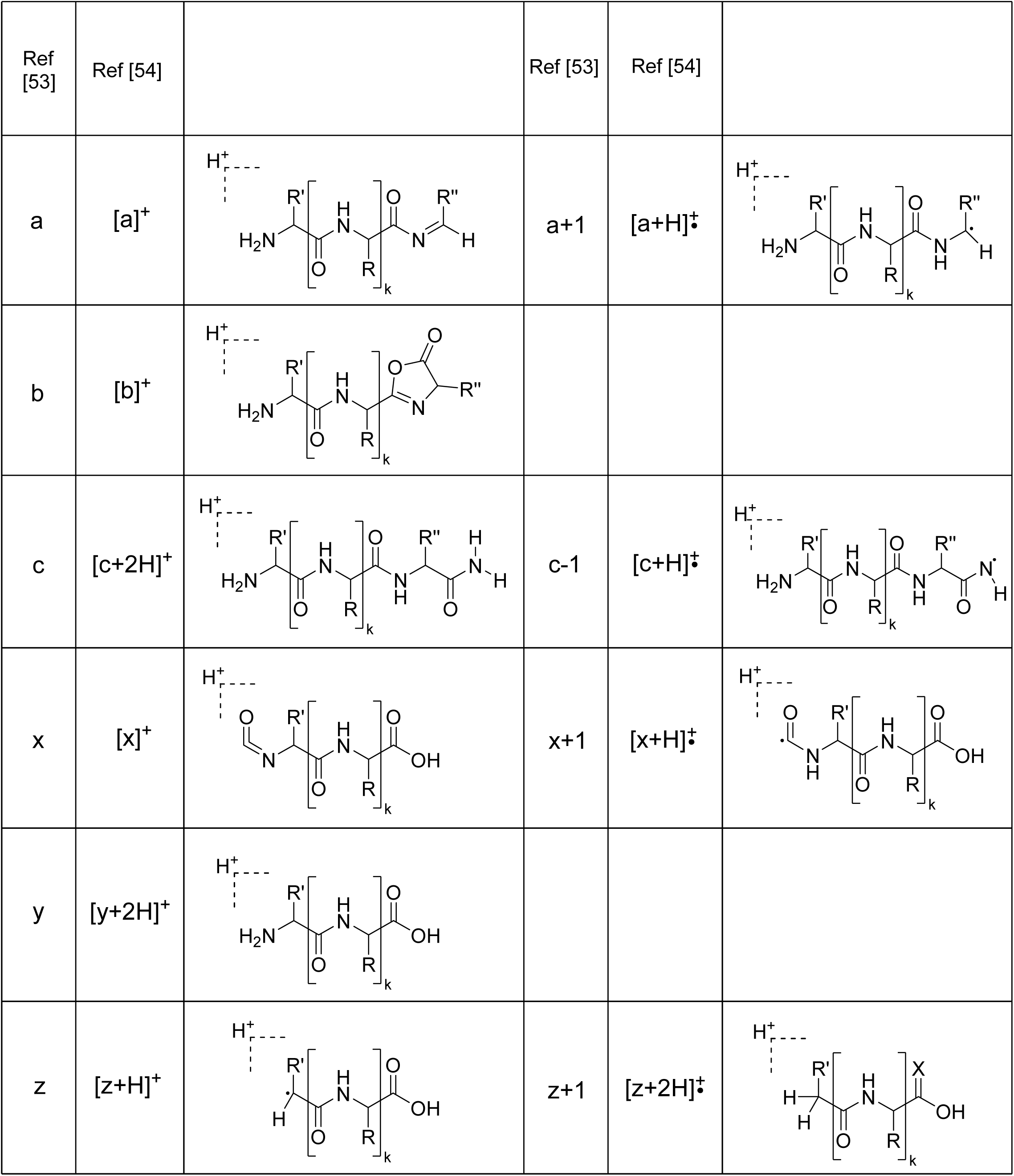
Definitions of ion types. Ion nomenclature proposed by Biemann (Ref [53], used in this work) and Chu *et al.* (Ref [54]) and corresponding elemental compositions of fragments

**Supplementary Figure S2.**
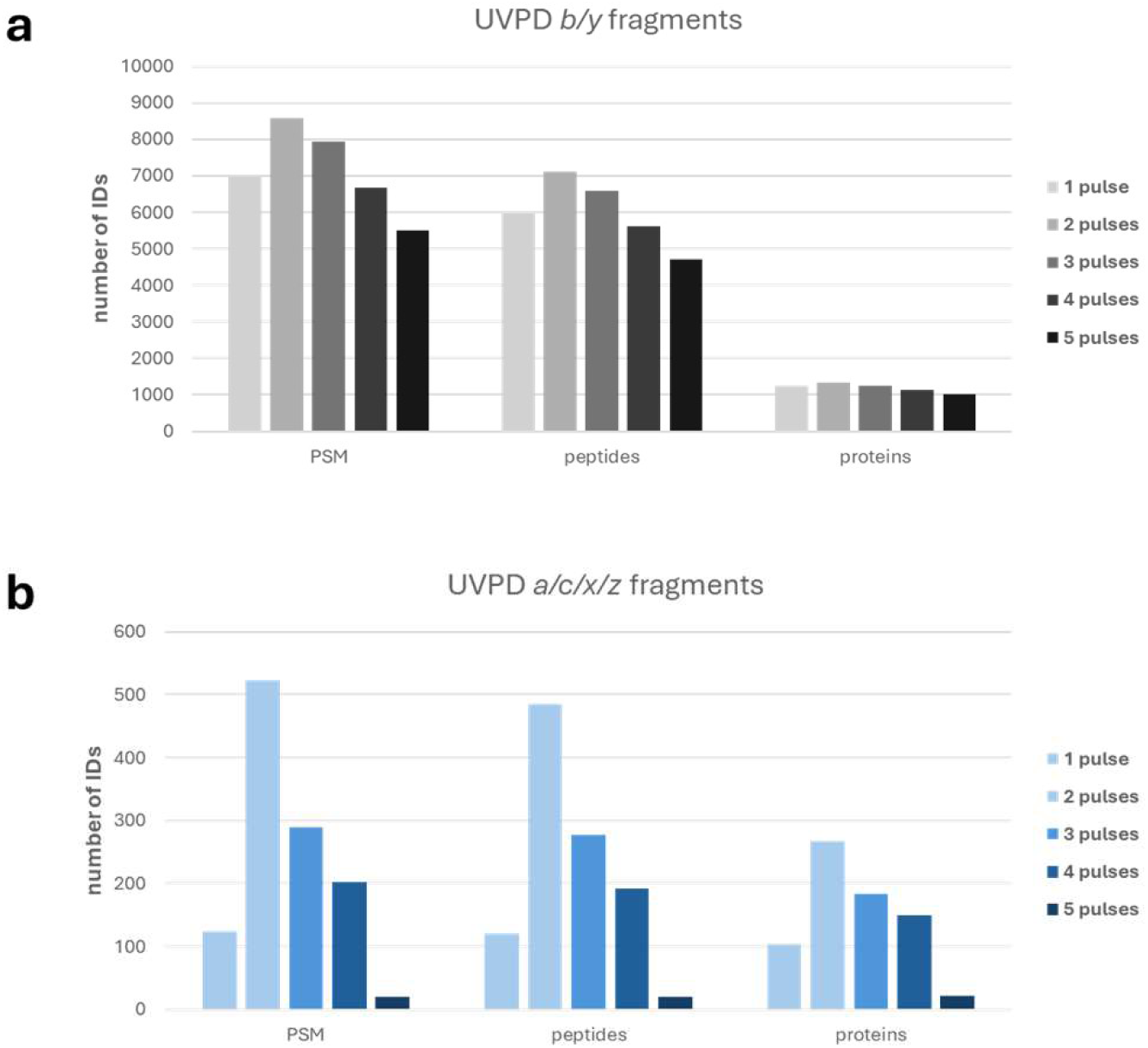
Initial optimization of UVPD. Numbers of peptide-spectrum matches (PSMs), peptides and proteins identified in the UVPD LCMS experiments using *b* and *y* fragments (a) or *a,c,x,z* fragments (b) for data analysis. 30 min LC gradients were used for the analysis of tryptic digest of human cells.

**Supplementary Figure S3.**
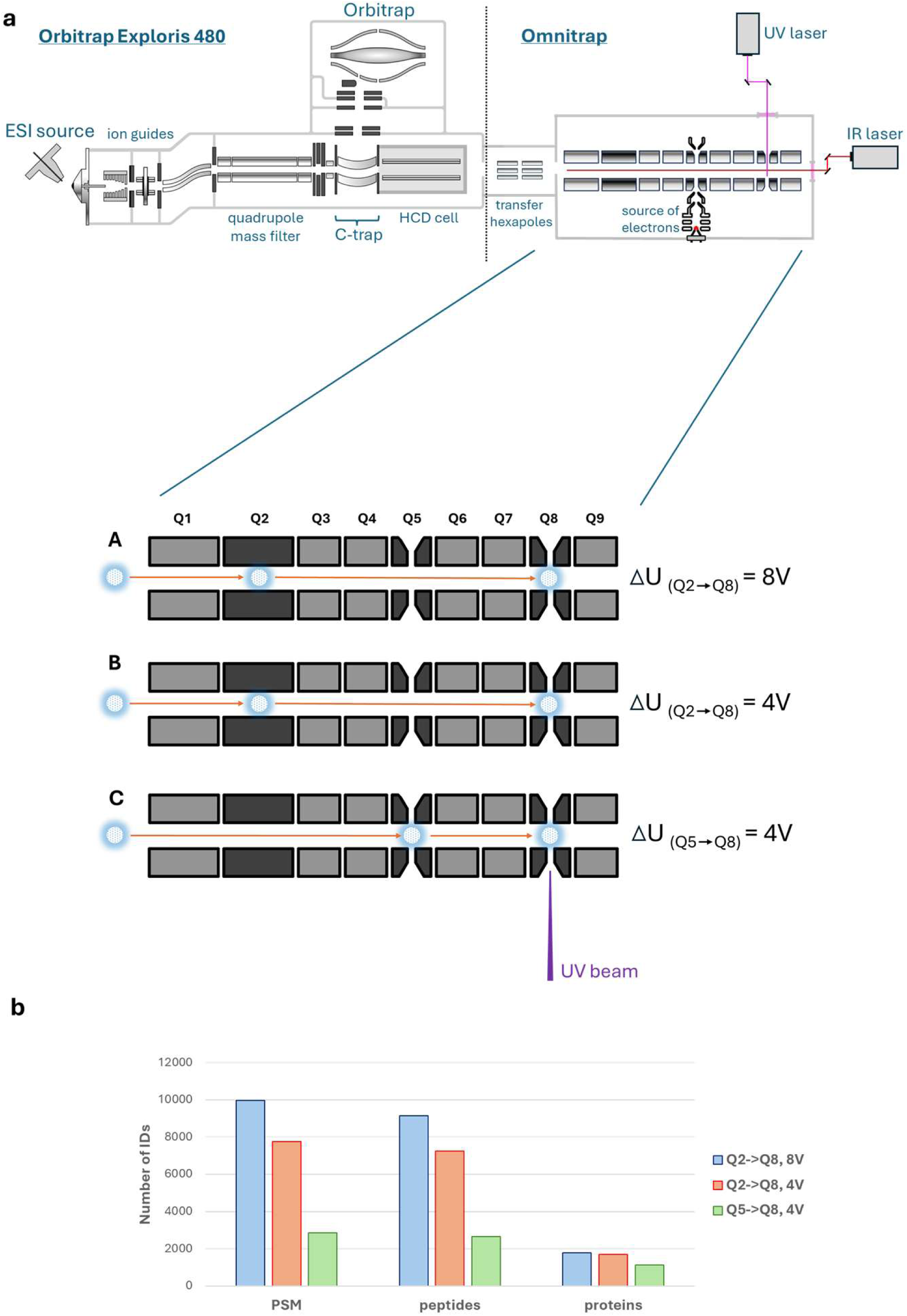
Ion transfer designs for UVPD and their effect on collisional background. a) Schematic representations of different designs of ion transfer to Q8 segment of the Omnitrap; b) numbers of peptide-spectrum matches (PSMs), peptides, and proteins identified in the LCMS experiments using three different designs of ions transfer without UV laser triggering. *b* and *y* fragments were used for data analysis.

**Supplementary Figure S4.**
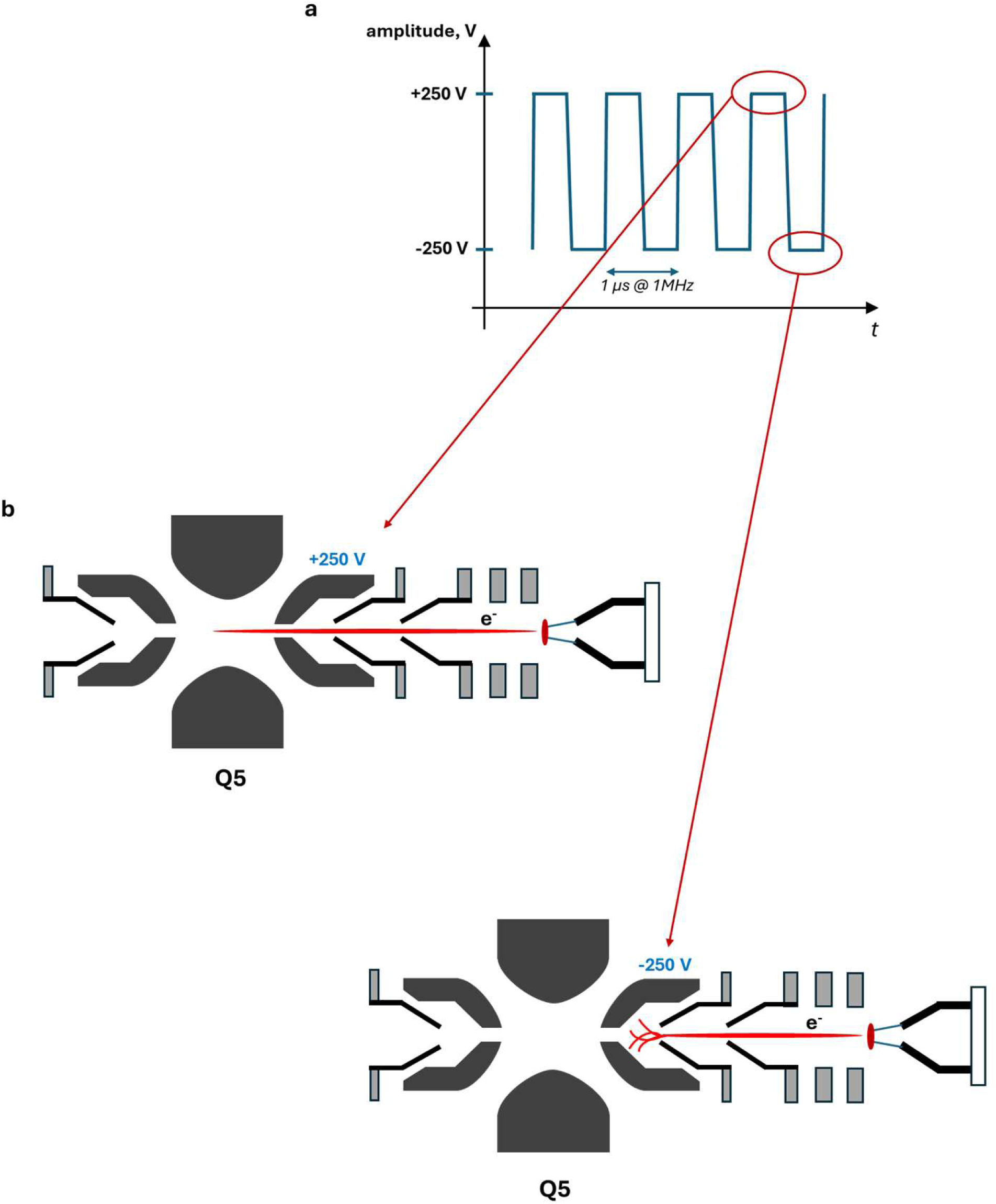
RF potentials and ejection of electrons into Q5 segment of the Omnitrap. **a**, Time scan of the electric potential at Q5 electrode, where the source of electrons is installed; **b,** schematic of the injection of electrons into the Q5 segment of the Omnitrap during positive (top) and negative (bottom) RF phases.

**Supplementary Figure S5.**
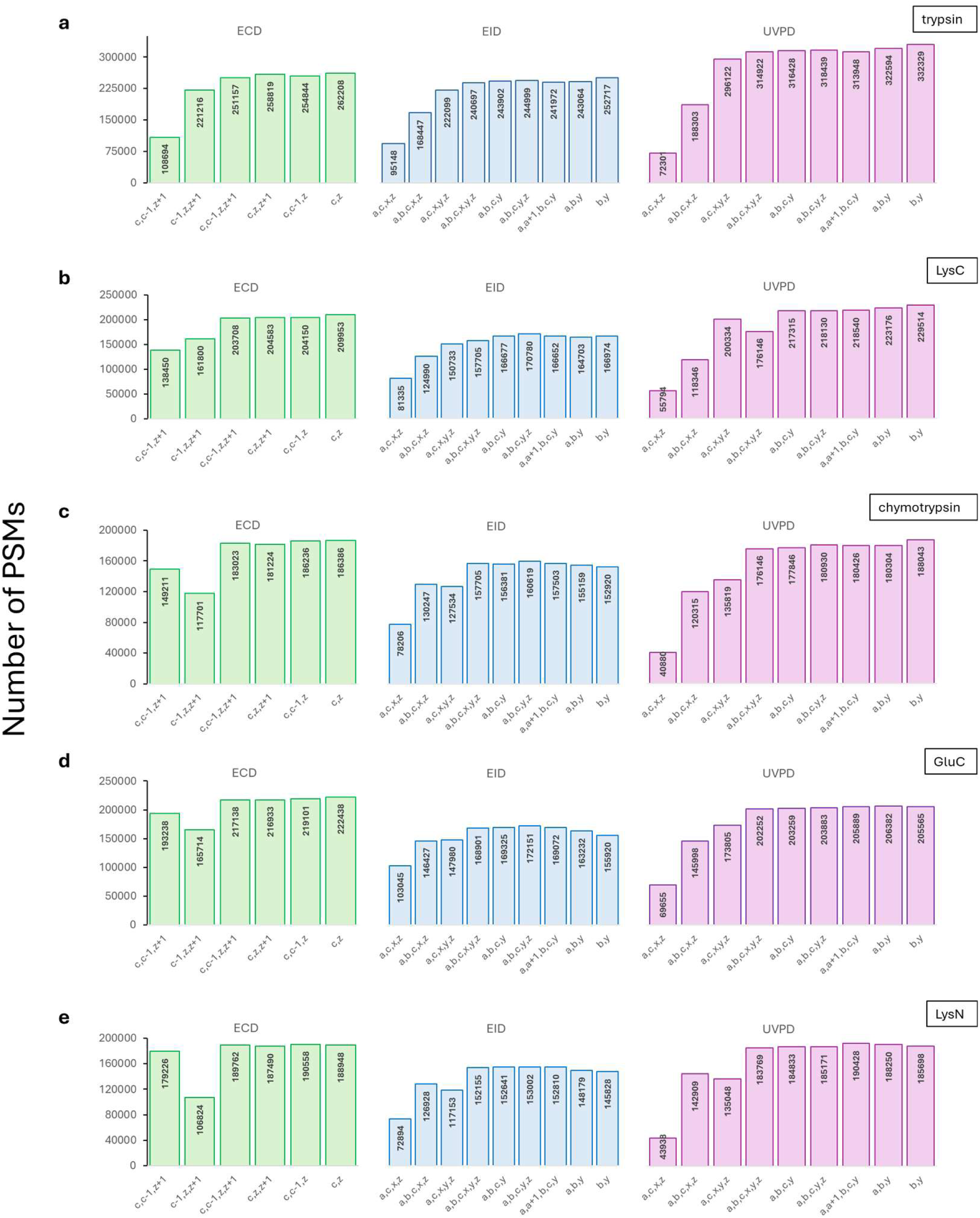
Breakdown of the contribution of each enzyme to the total numbers of PSMs. Numbers of peptide-spectrum matches (PSMs) in ECD, EID, and UVPD experiments identified using different combinations of fragment types in trypsin (a), LysC (b), chymotrypsin (c), GluC (d), and LysN (e) datasets.

**Supplementary Figure S6.**
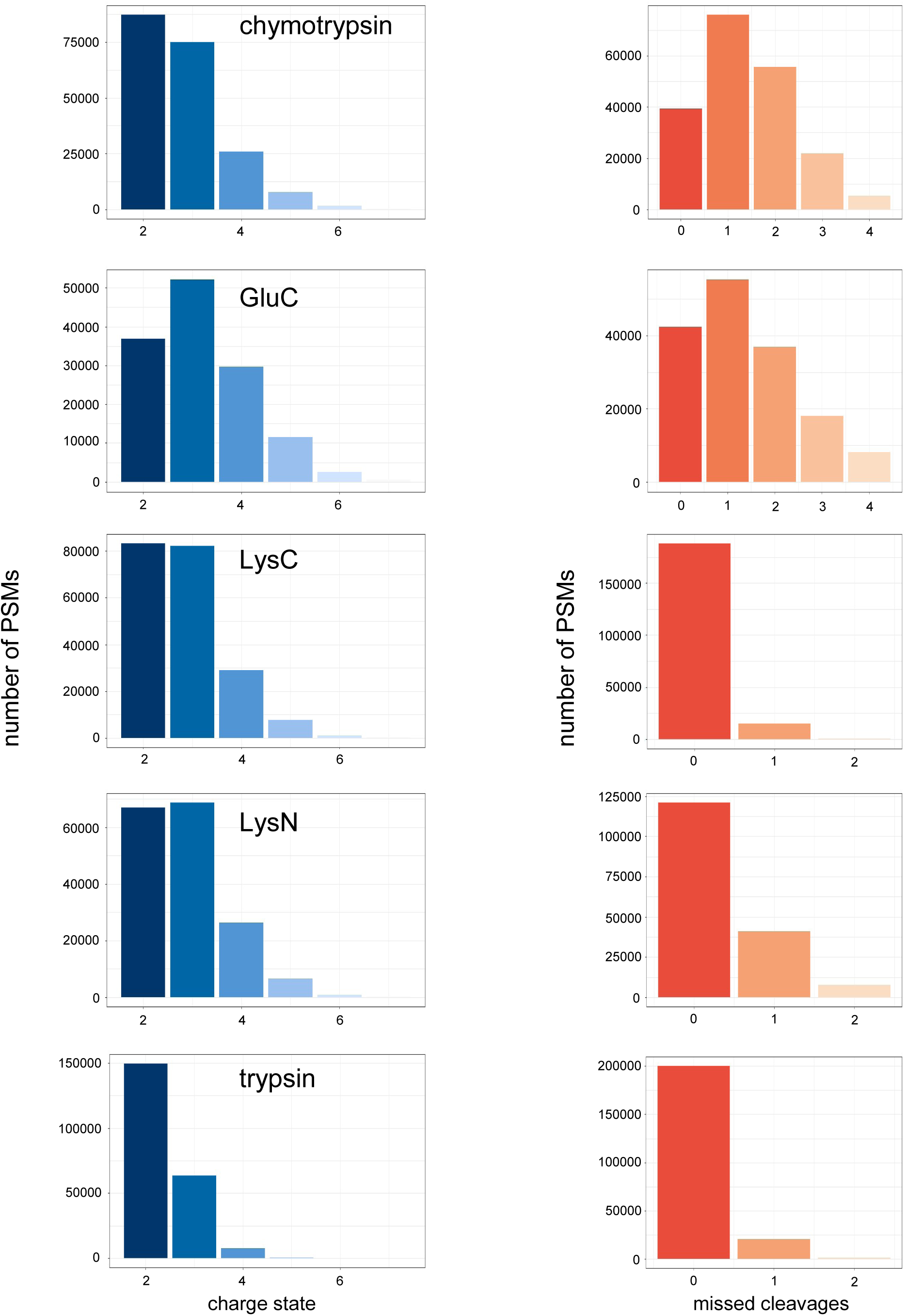
Enzyme-specific physicochemical properties of UVPD precursors. Per-enzyme distributions of charge states (left) and missed cleavages (right) for all PSMs identified in 2D-LCMS-UVPD experiments.

**Supplementary Figure S7.**
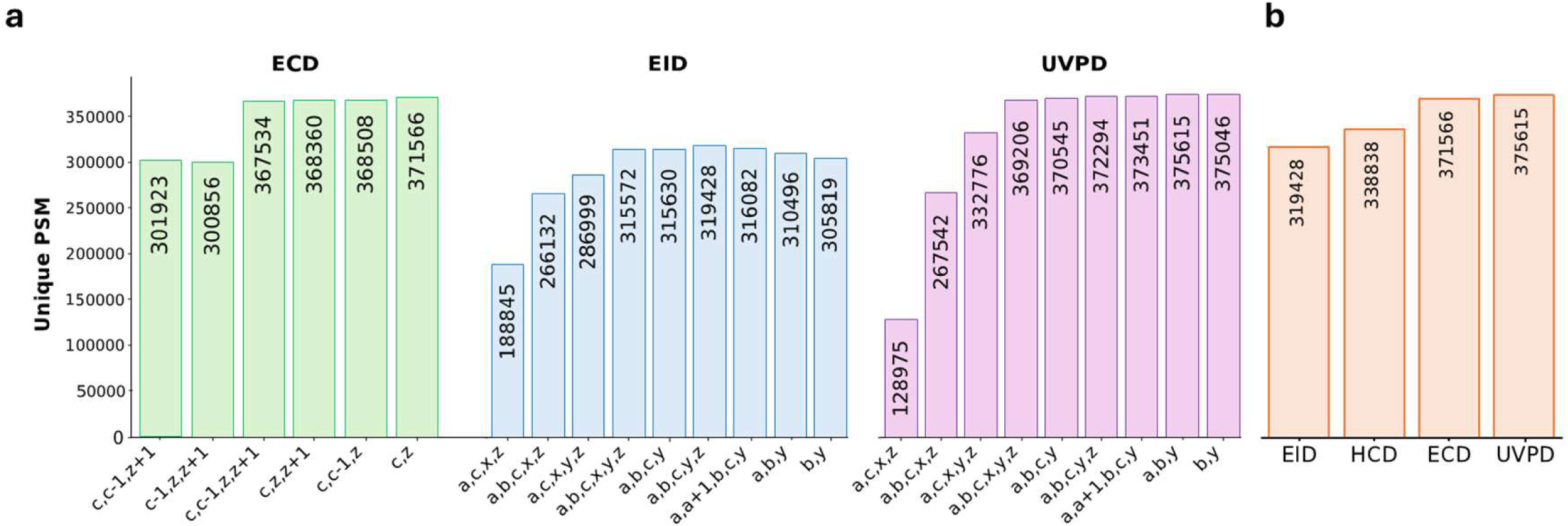
Unique PSMs in ECD, EID, and UVPD data. **a**, Total numbers of unique peptide-spectrum matches (unique combination of amino-acid sequence, charge, and modification) identified using different combinations of fragment types in ECD, EID, and UVPD experiments; **b,** highest number of PSMs from (a) together with the HCD data.

**Supplementary Figure S8.**
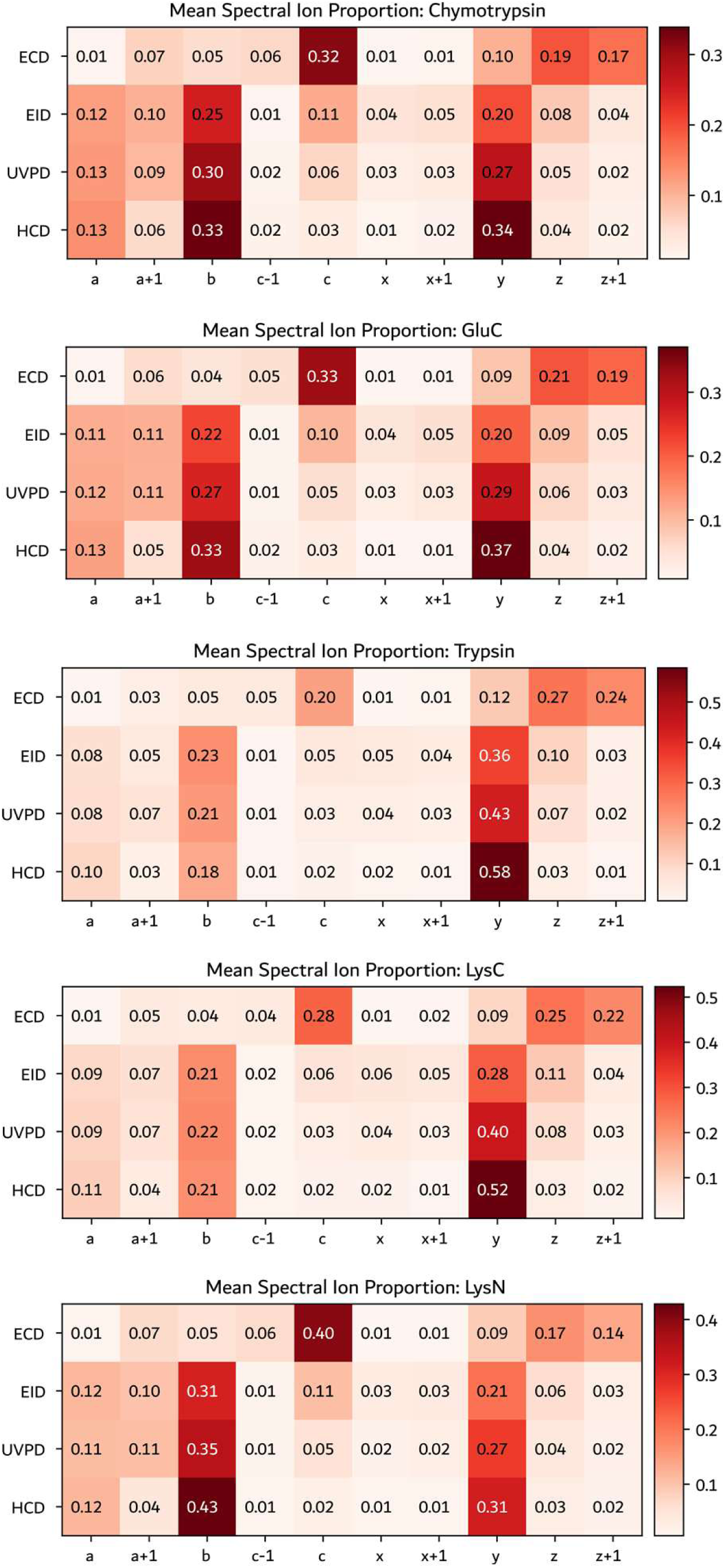
Averaged spectral annotations per enzyme. Heatmap of average proportions of fragment ions of different types among all annotated ions in ECD, EID, HCD, and UVPD spectra plotted per enzyme. Annotation was performed for 10 ion types: *a*, *a+1*, *b*, *c-1*, *c*, *x*, *x+1*, *y*, *z*, *z+1*.

**Supplementary Figure S9.**
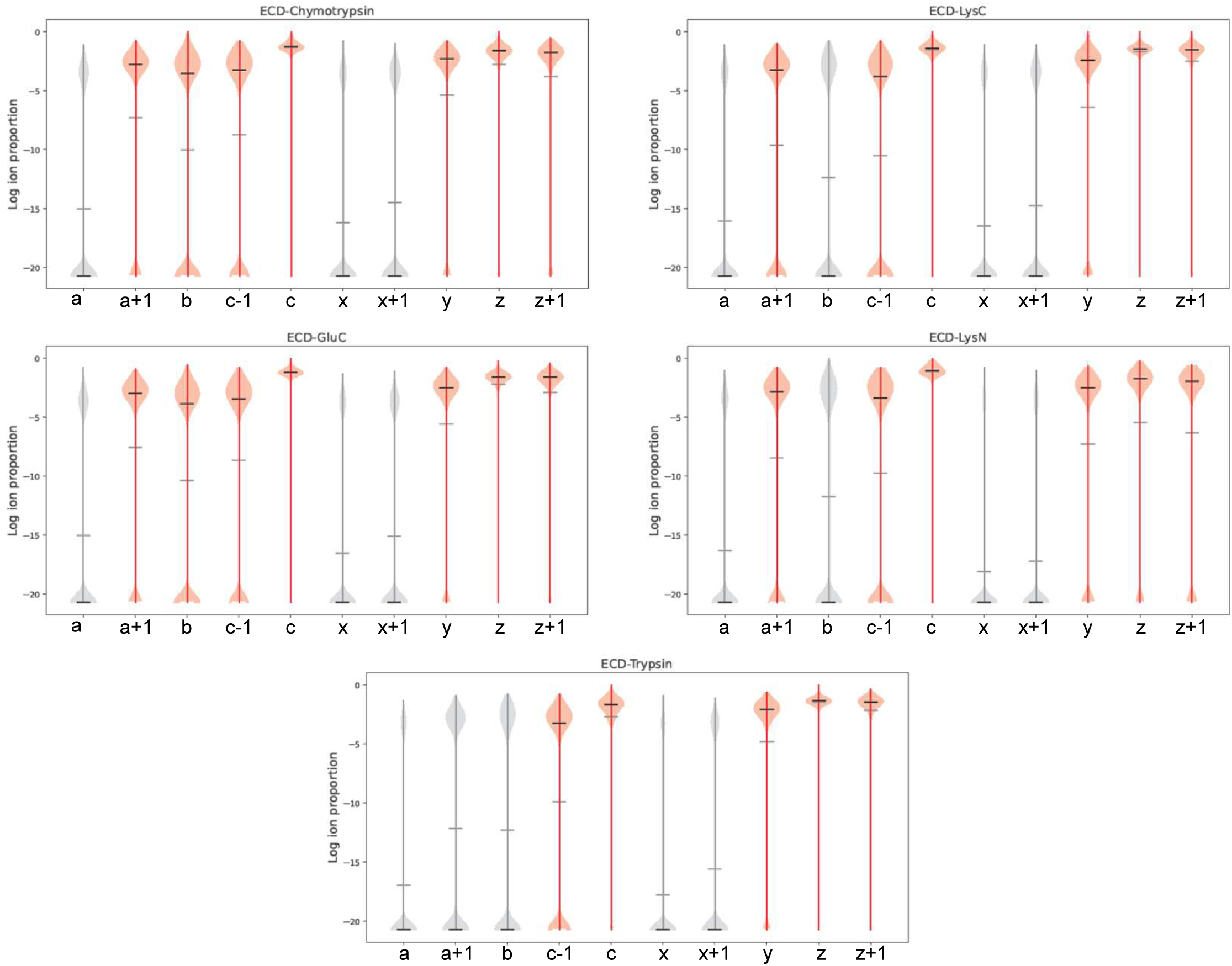
Log ion-proportions of ions annotated in ECD data plotted per enzyme. Ions annotated with negligible frequencies (<4% of all annotated ions) are greyed out. Horizontal grey and black lines correspond to the median and mean values, respectively.

**Supplementary Figure S10.**
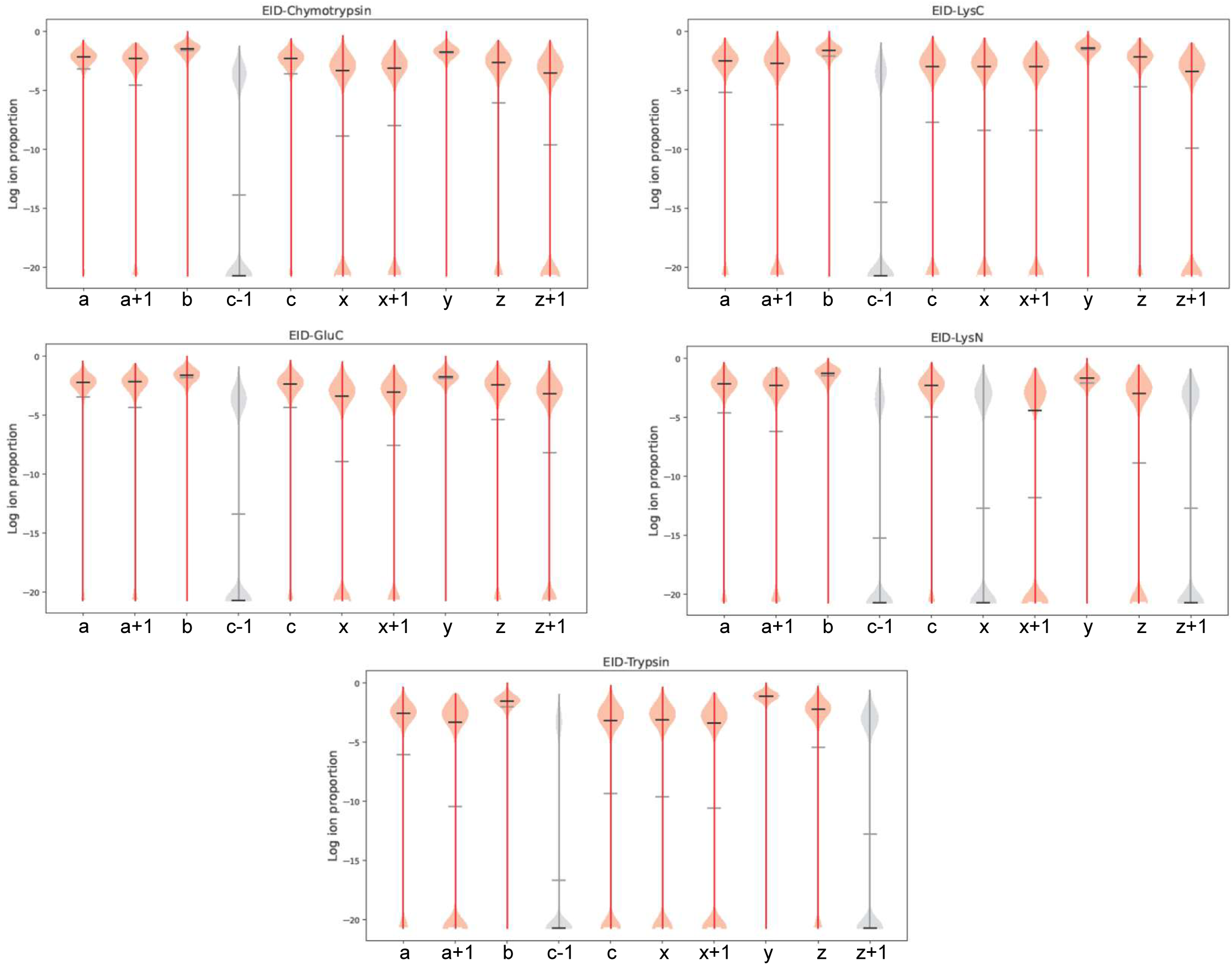
Log ion-proportions of ions annotated in EID data plotted per enzyme. Ions annotated with negligible frequencies (<4% of all annotated ions) are greyed out. Horizontal grey and black lines correspond to the median and mean values, respectively.

**Supplementary Figure S11.**
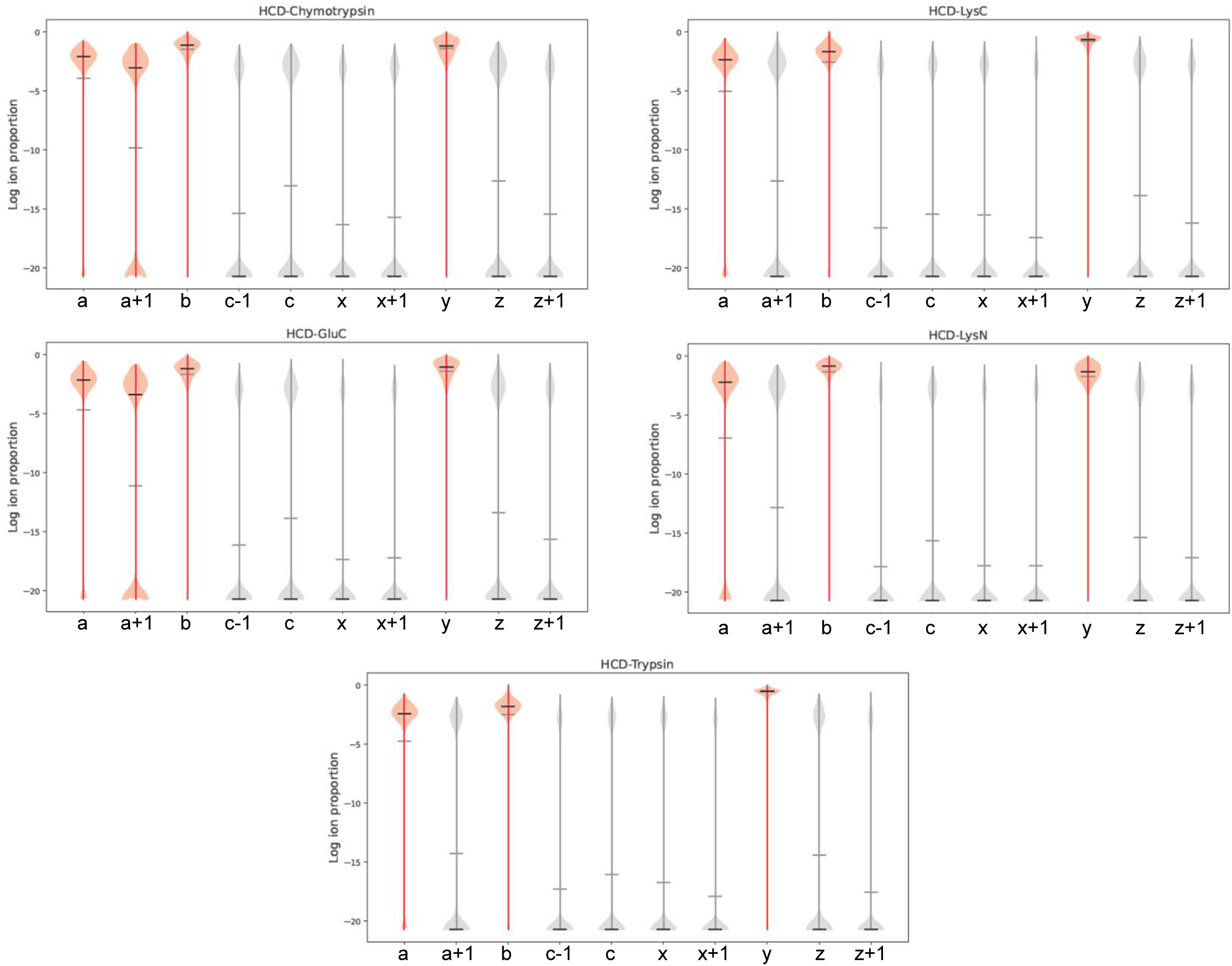
Log ion-proportions of ions annotated in HCD data plotted per enzyme. Ions annotated with negligible frequencies (<4% of all annotated ions) are greyed out. Horizontal grey and black lines correspond to the median and mean values, respectively.

**Supplementary Figure S12.**
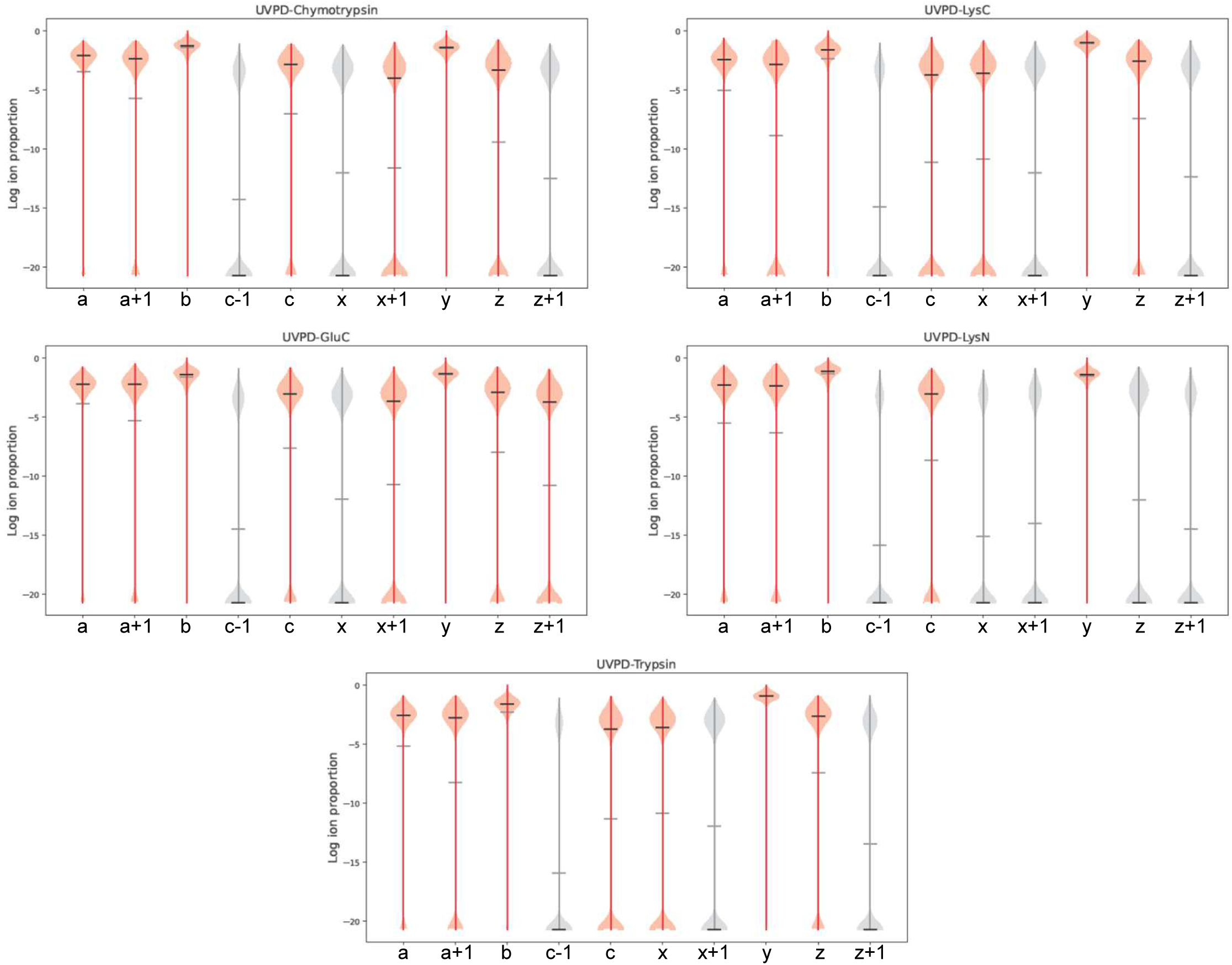
Log ion-proportions of ions annotated in UVPD data plotted per enzyme. Ions annotated with negligible frequencies (<4% of all annotated ions) are greyed out. Horizontal grey and black lines correspond to the median and mean values, respectively.

**Supplementary Figure S13.**
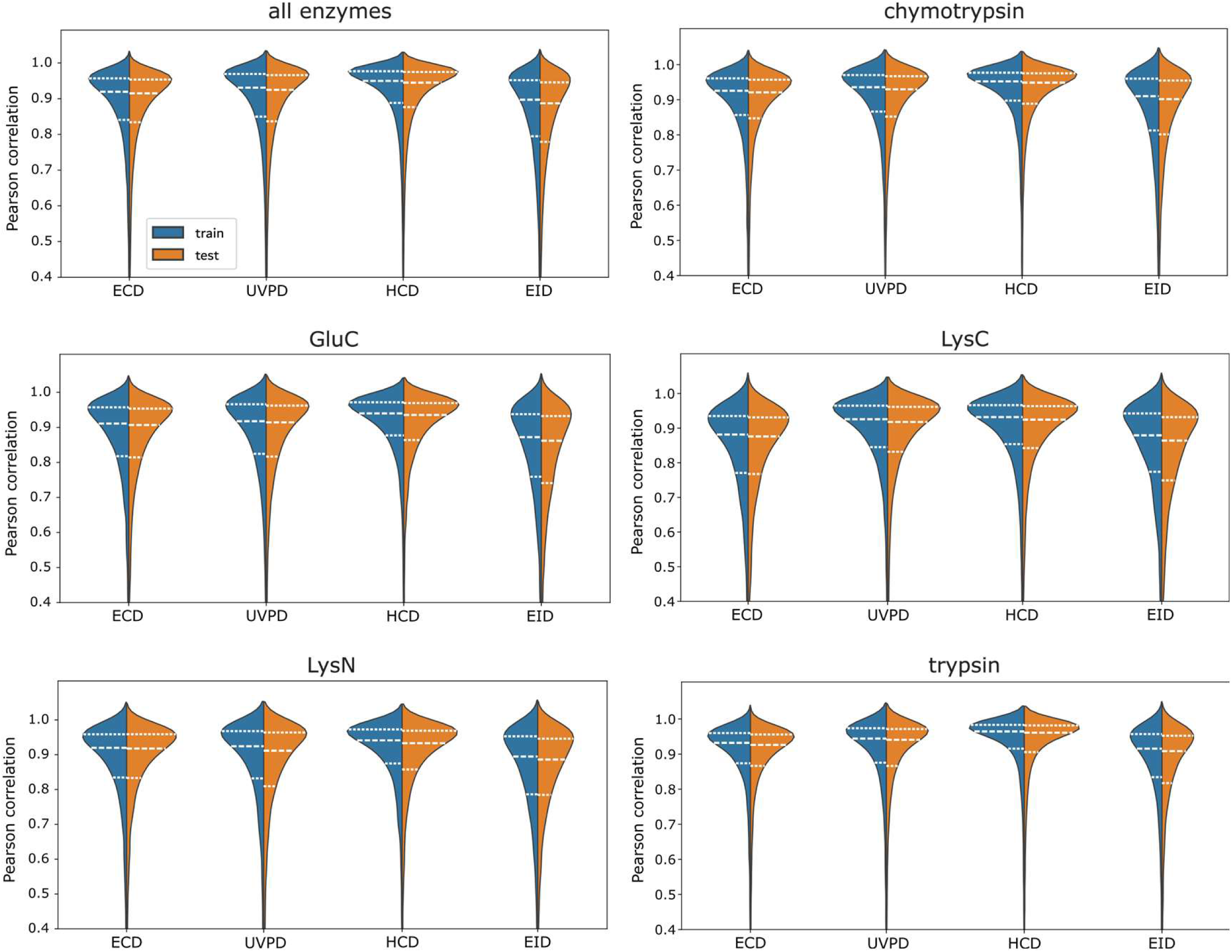
Pearson correlations per enzyme per fragmentation technique. Distributions of Pearson correlation between experimental and predicted fragmentation spectra plotted for train and test datasets per enzyme per fragmentation method. Horizontal dashed lines correspond to 25, 50, and 75% percentiles. Distributions protruding beyond 1.0 are plotting artefacts.

**Supplementary Figure S14.**
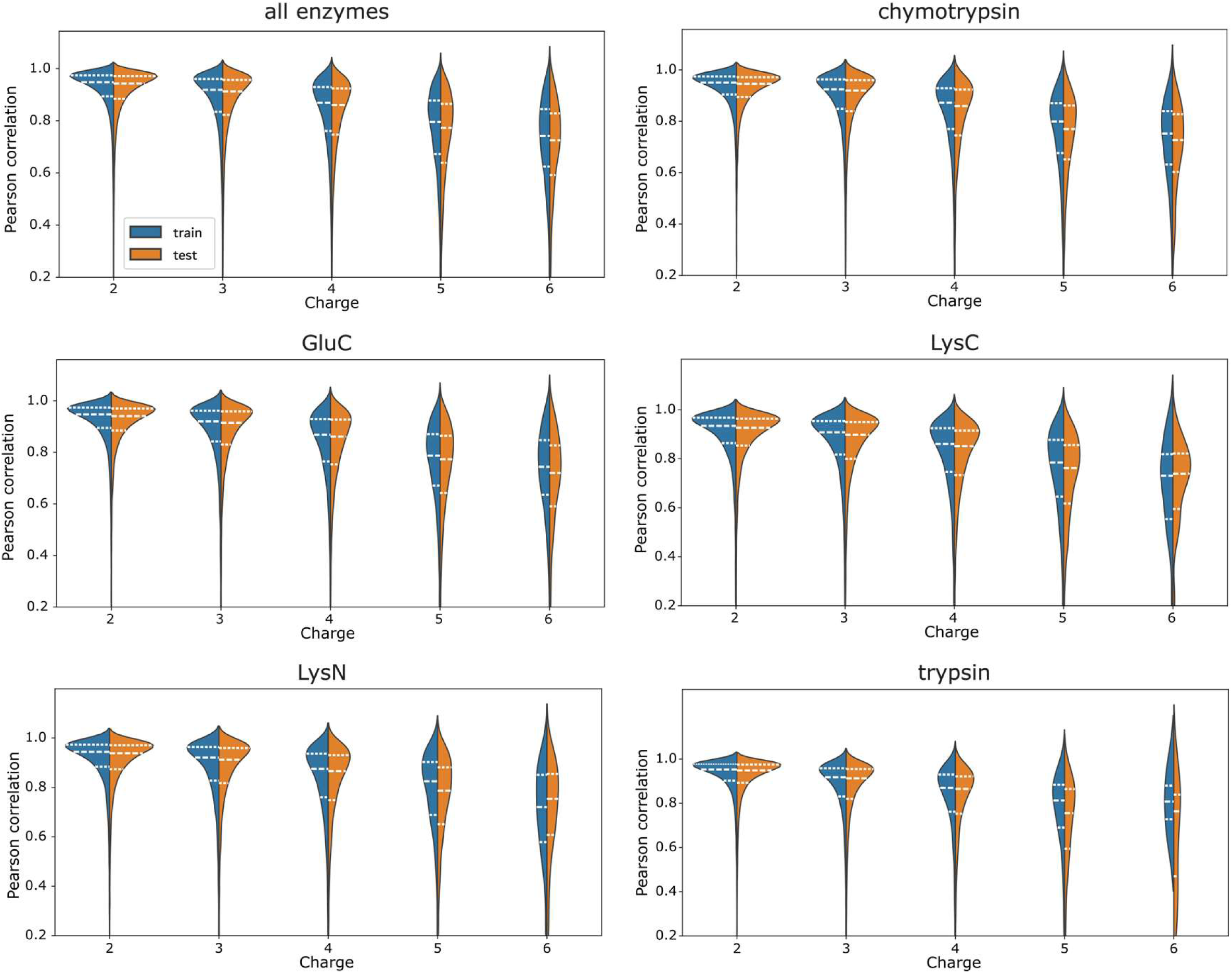
Pearson correlations per enzyme per charge state. Distributions of Pearson correlation between experimental and predicted fragmentation spectra plotted for train and test datasets per enzyme per charge state. Horizontal dashed lines correspond to 25, 50, and 75% percentiles. Distributions protruding beyond 1.0 are plotting artefacts.

**Supplementary Figure S15.**
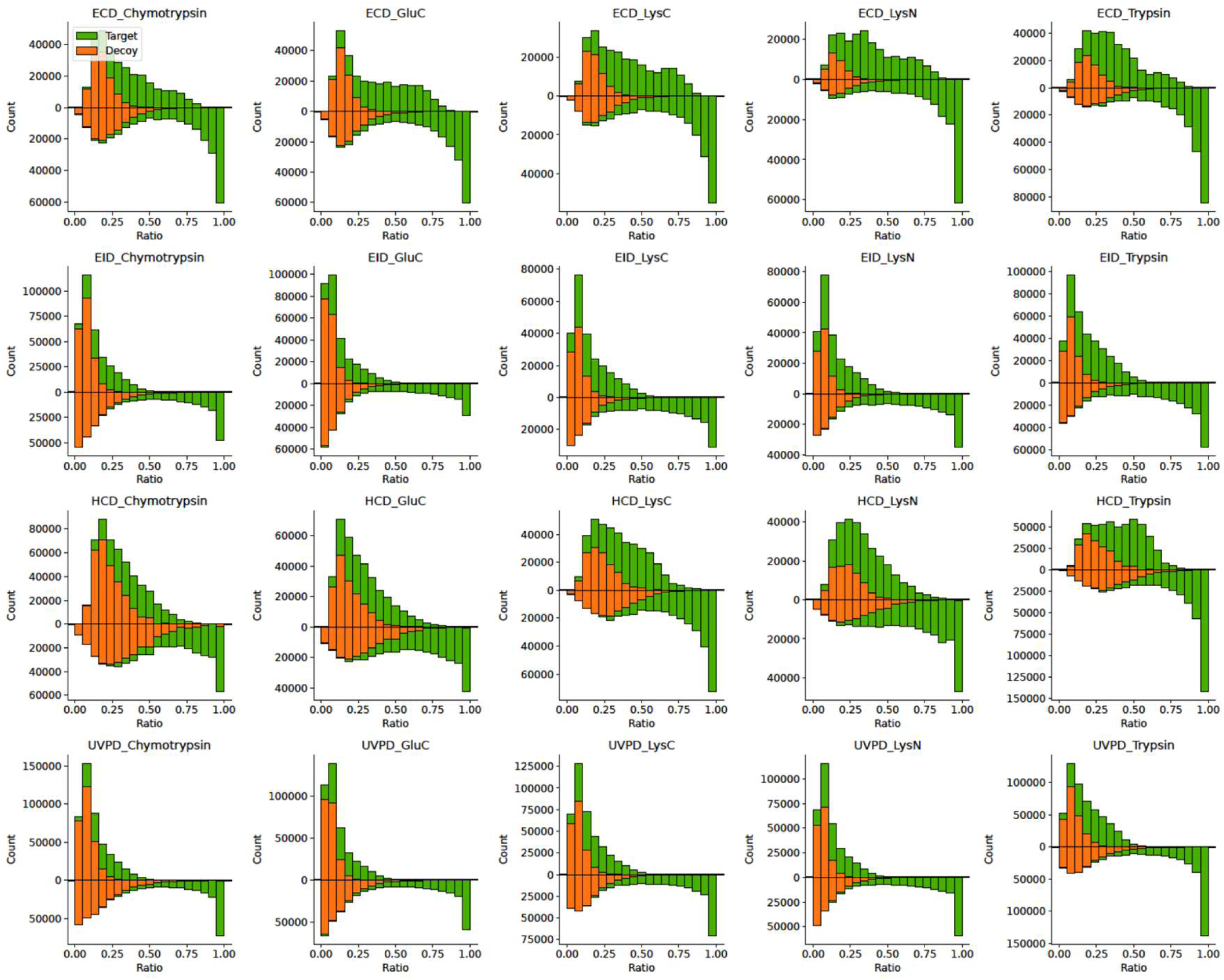
Predictions of fragment’s presence or absence in a spectrum help separate target PSMs from decoy. Histograms of the ratios of experimentally observed ions to all theoretically possible fragment (upper distributions); Histograms of the ratios of experimentally observed and predicted ions to all predicted ions (bottom distributions).

**Supplementary Figure S16.**
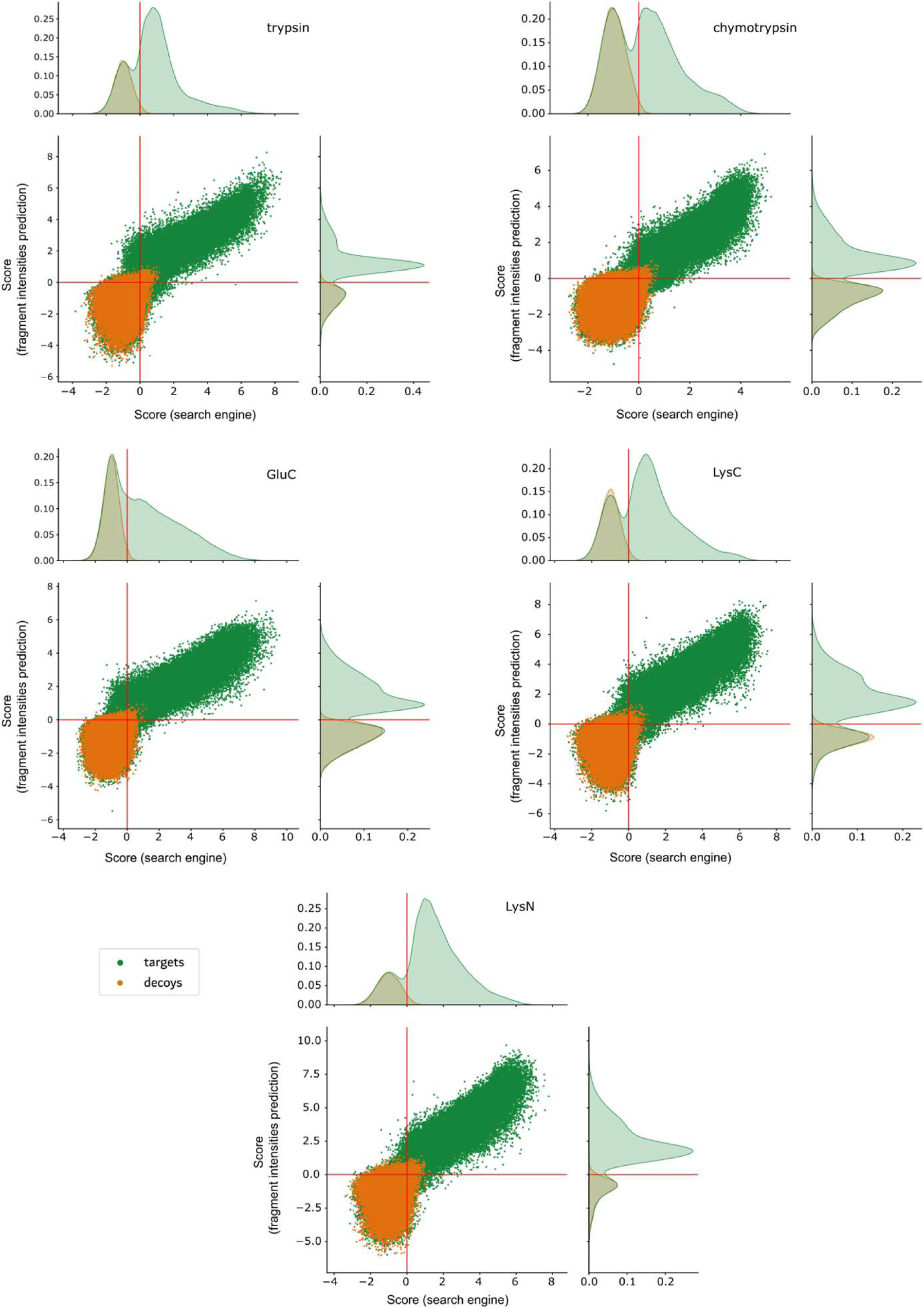
Oktoberfest rescoring separates target from decoy. Correlation of Percolator scores for all target (green) and decoy (orange) PSMs in ECD data acquired from rescoring the MSFragger (top) and Oktoberfest (right) sets of scores plotted per enzyme. The red solid lines indicate the 1% PSM level FDR cutoffs in MSFragger and Oktoberfest score distributions.

**Supplementary Figure S17.**
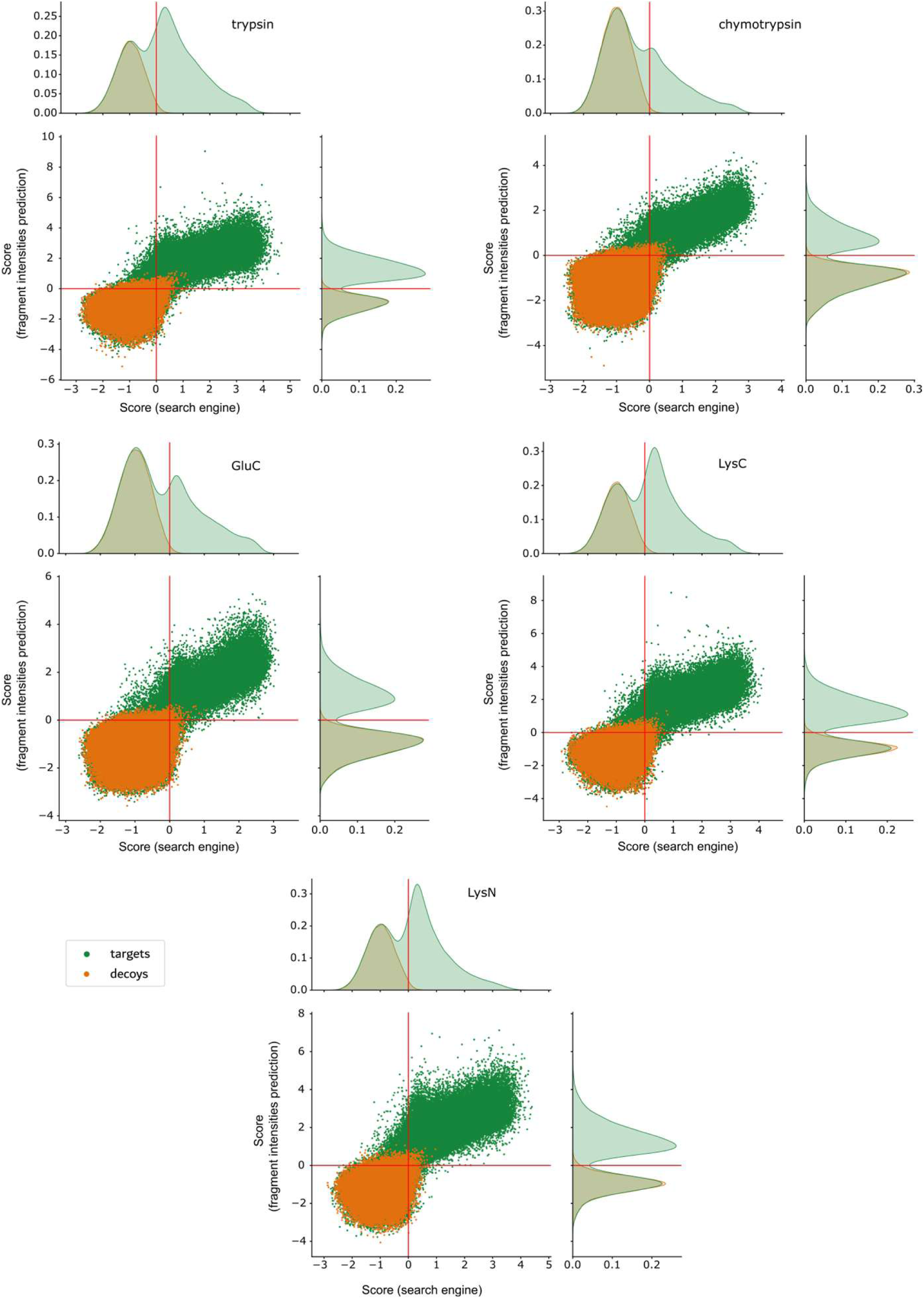
Oktoberfest rescoring separates target from decoy. Correlation of Percolator scores for all target (green) and decoy (orange) PSMs in EID data acquired from rescoring the MSFragger (top) and Oktoberfest (right) sets of scores plotted per enzyme. The red solid lines indicate the 1% PSM level FDR cutoffs in MSFragger and Oktoberfest score distributions.

**Supplementary Figure S18.**
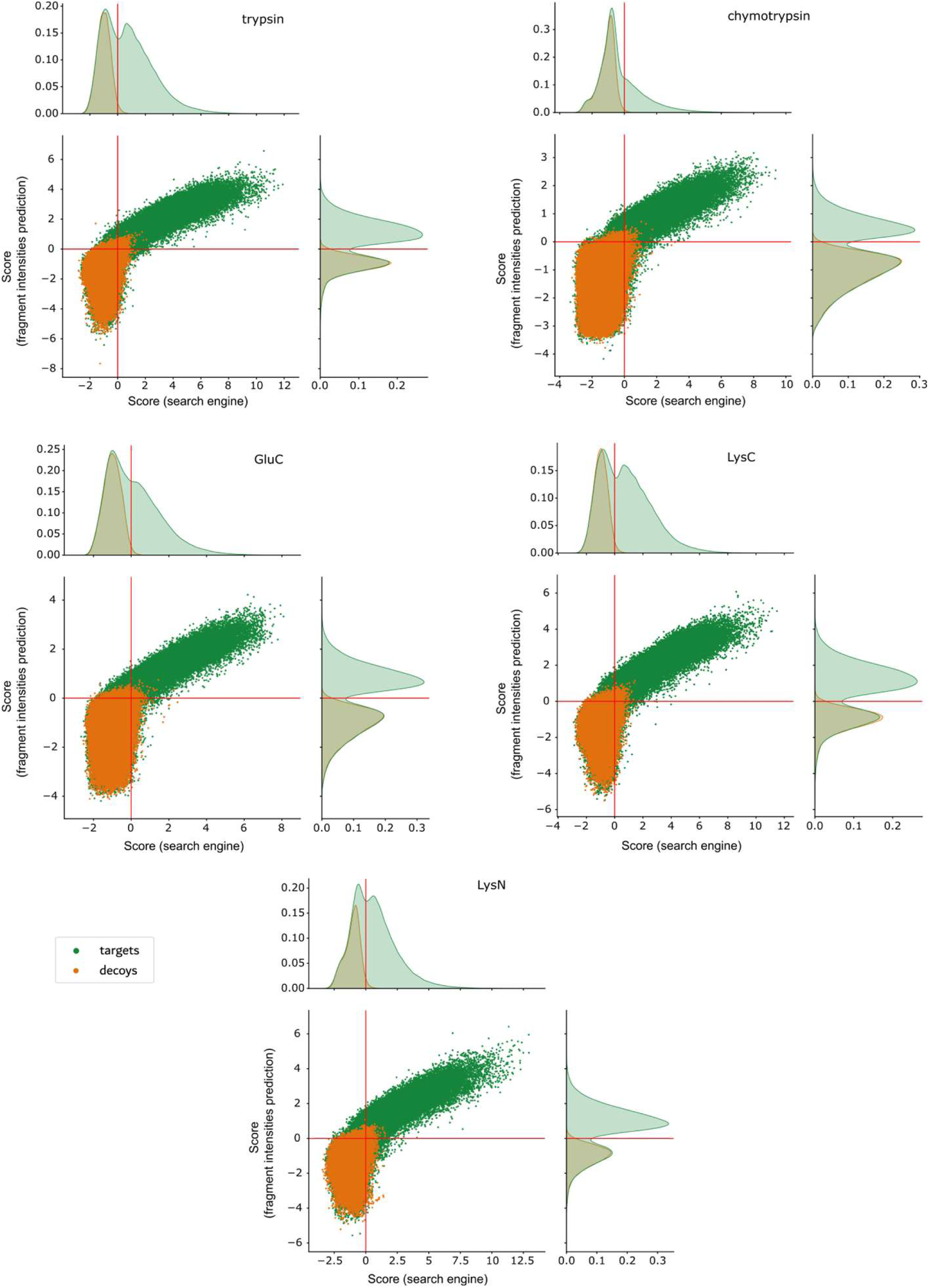
Oktoberfest rescoring separates target from decoy. Correlation of Percolator scores for all target (green) and decoy (orange) PSMs in HCD data acquired from rescoring the MSFragger (top) and Oktoberfest (right) sets of scores plotted per enzyme. The red solid lines indicate the 1% PSM level FDR cutoffs in MSFragger and Oktoberfest score distributions.

**Supplementary Figure S19.**
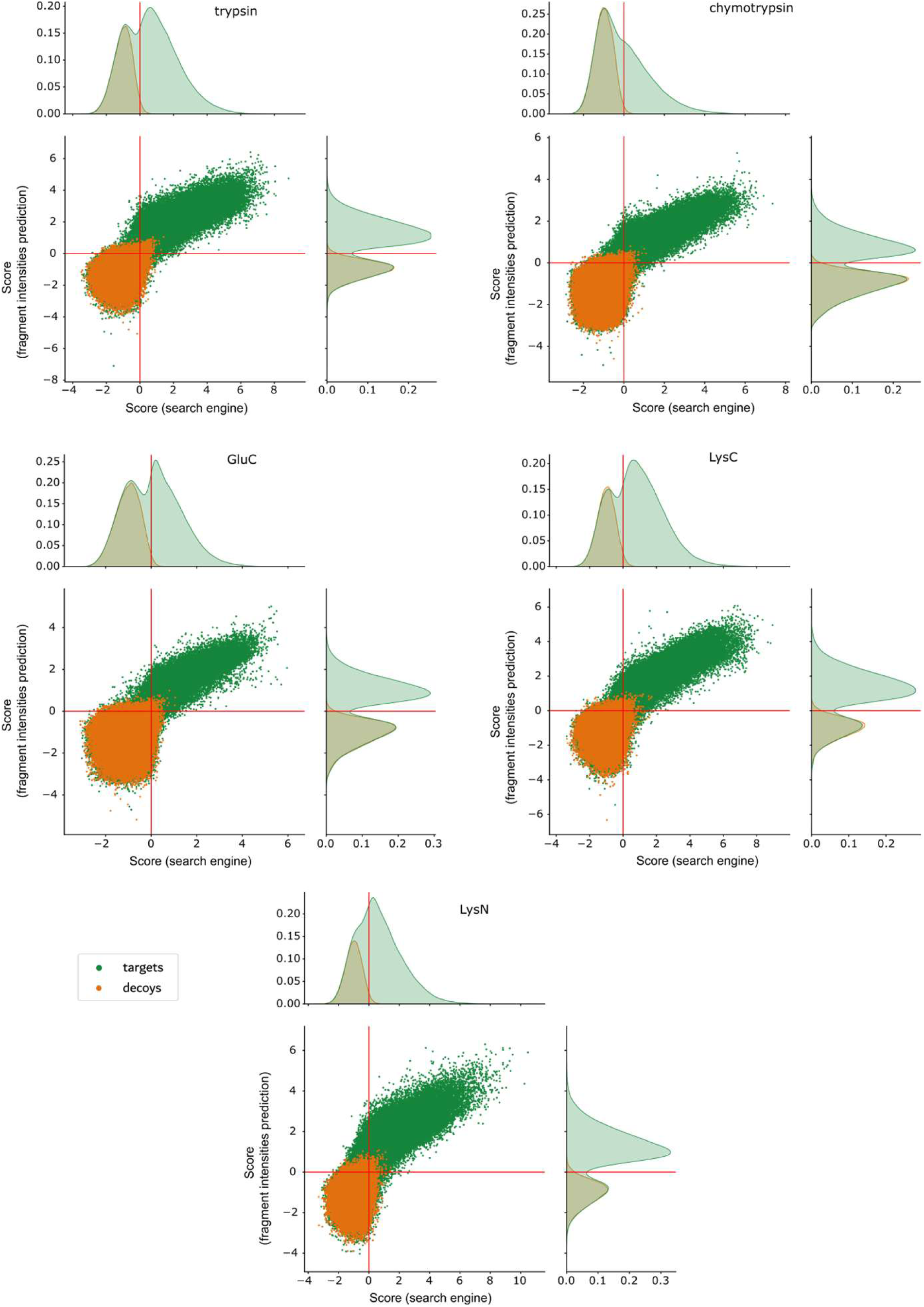
Oktoberfest rescoring separates target from decoy. Correlation of Percolator scores for all target (green) and decoy (orange) PSMs in UVPD data acquired from rescoring the MSFragger (top) and Oktoberfest (right) sets of scores plotted per enzyme. The red solid lines indicate the 1% PSM level FDR cutoffs in MSFragger and Oktoberfest score distributions.

**Supplementary Figure S20.**
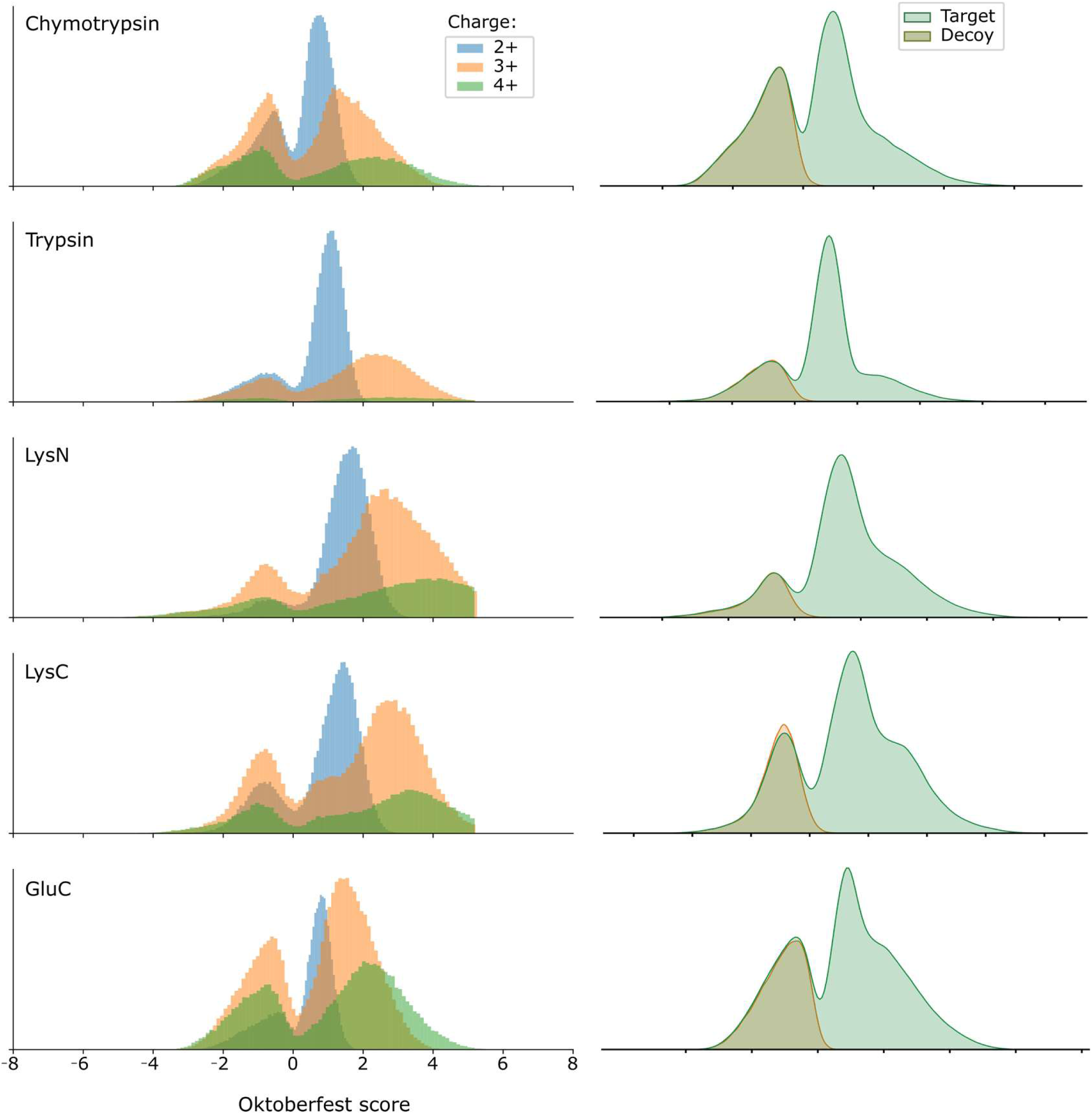
Distributions of target PSMs in ECD are charge-separated after rescoring in Oktoberfest. **Left**: PSM-level distributions of Oktoberfest scores separated by charge state of precursors plotted for ECD data per enzyme. **Right**: the same distributions summed up and split between decoy and target (taken from Supplementary Fig. S16)

**Supplementary Figure S21.**
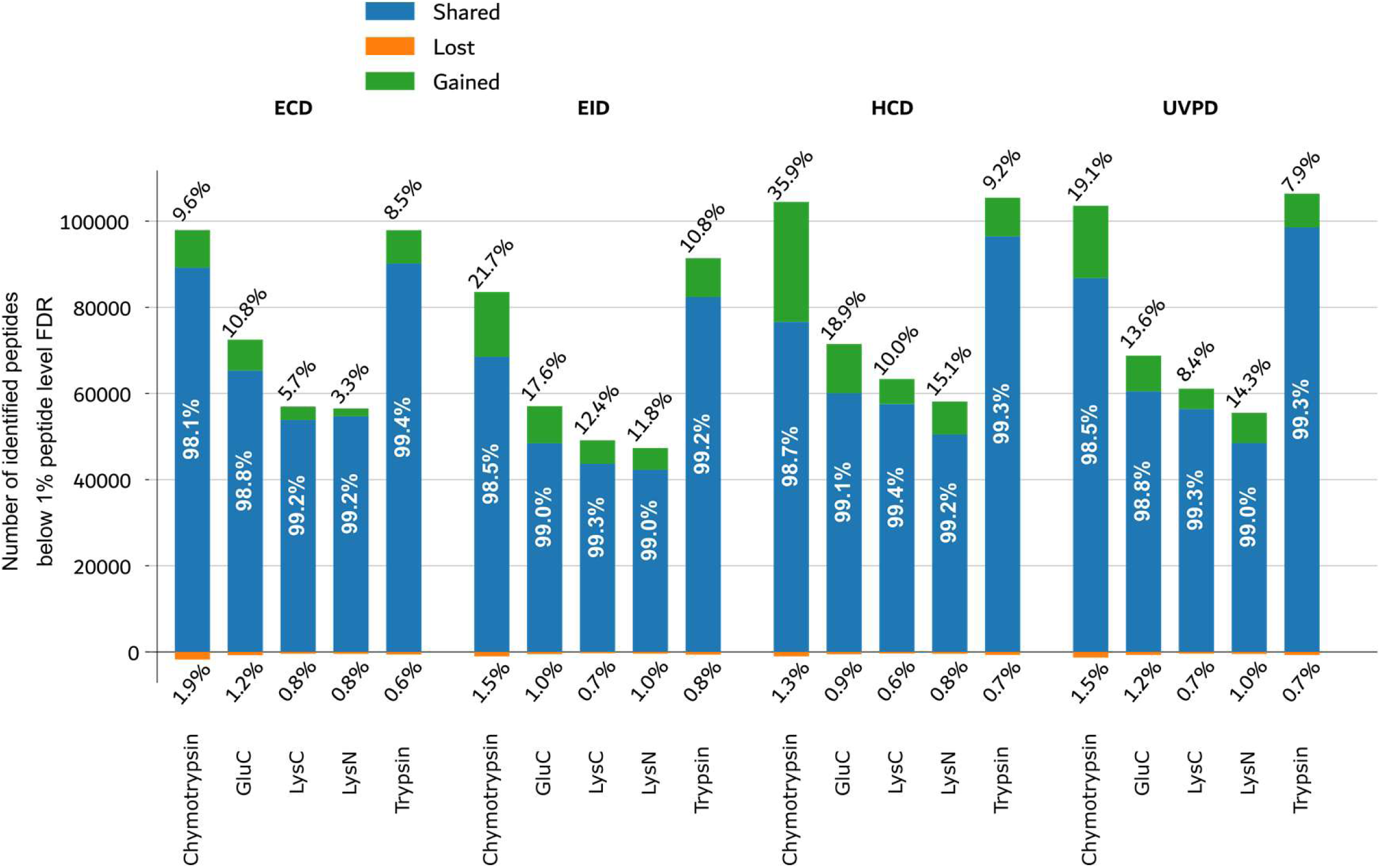
Gain/share/loss on the peptide level. Numbers of shared (blue), gained (green), and lost (orange) peptides identified at 1% peptide-level FDR using the Oktoberfest set of scores compared to original MSFragger search for each fragmentation technique per enzyme.

**Supplementary Figure S22.**
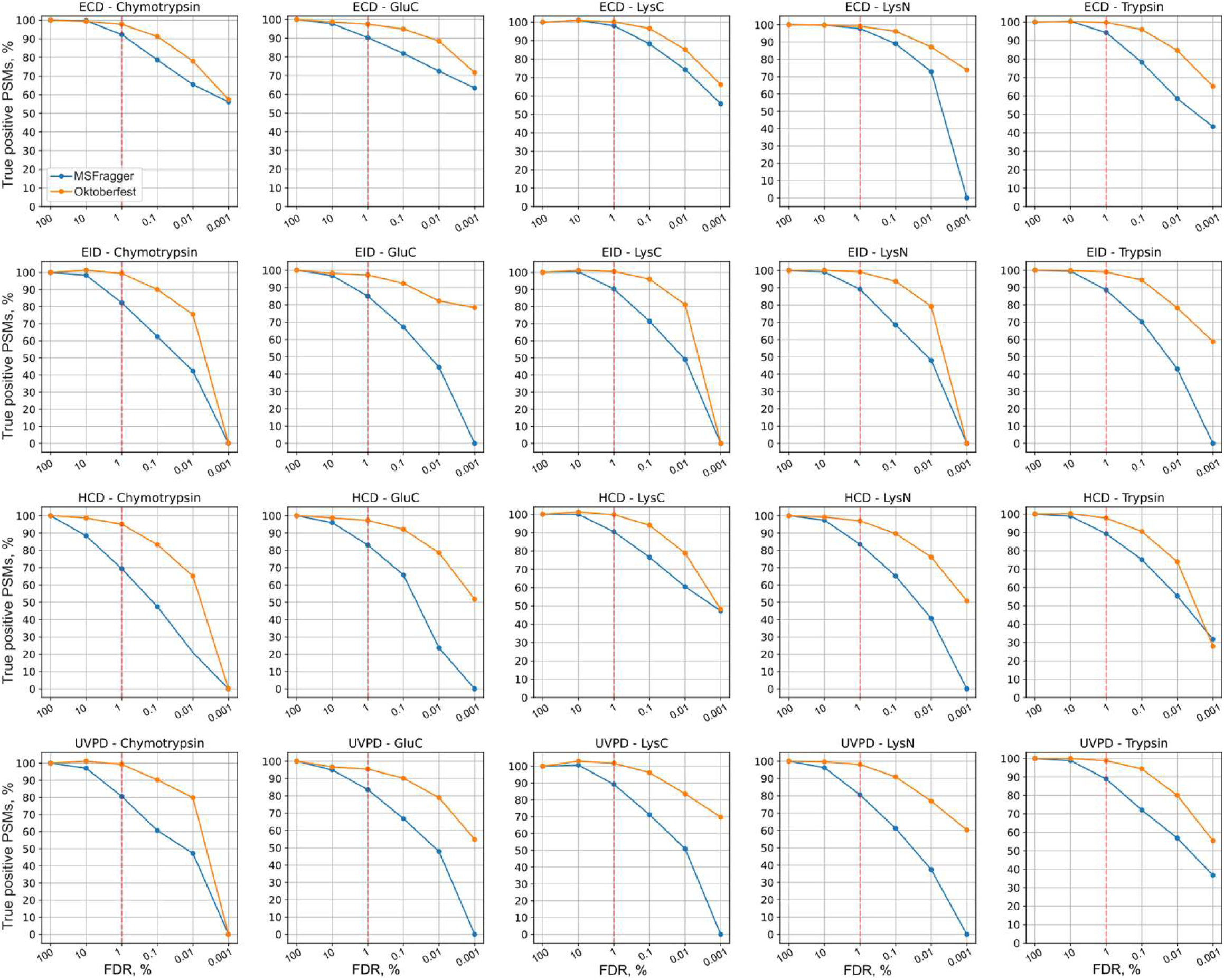
Identification of true positives at different FDR thresholds. Proportions of the numbers of true positive PSMs to the estimated maximum number of true positive PSMs acquired using original MSFragger and Oktoberfest scores at different values of PSM-level FDR for each fragmentation technique, per enzyme.

**Supplementary Figure S23.**
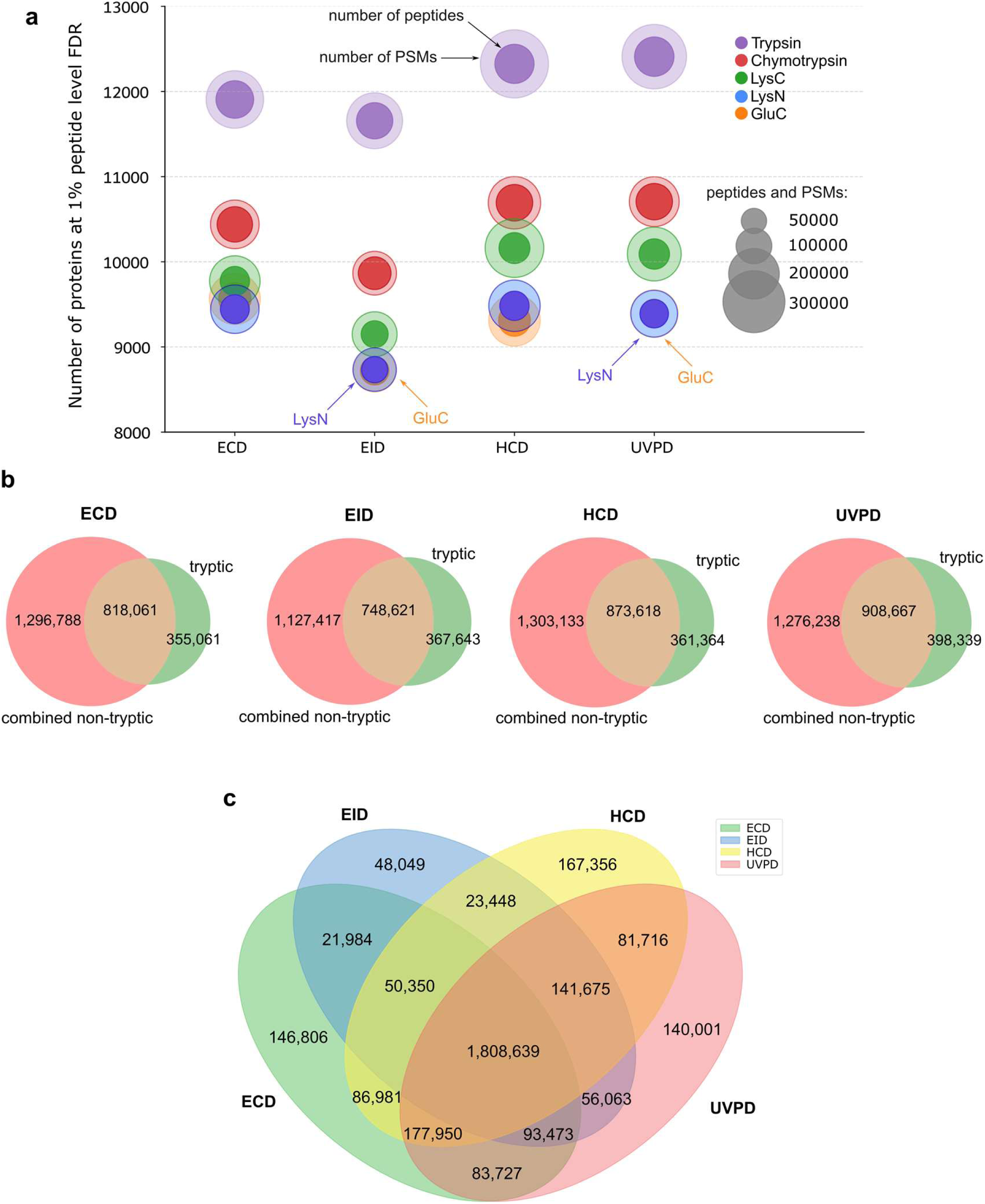
Relative performance of enzymes in DDA data. **a**, Numbers of PSMs (outer circles), peptides (inner circles) and proteins (y-axis) acquired using different fragmentation techniques and enzymes. **b,** Venn diagrams of total numbers of unique amino acids observed in tryptic datasets (green) and all non-tryptic datasets combined (red) using different fragmentation techniques. **c,** Venn diagram of unique amino acids observed using different fragmentation techniques across all enzymatic datasets.

**Supplementary Figure S24.**
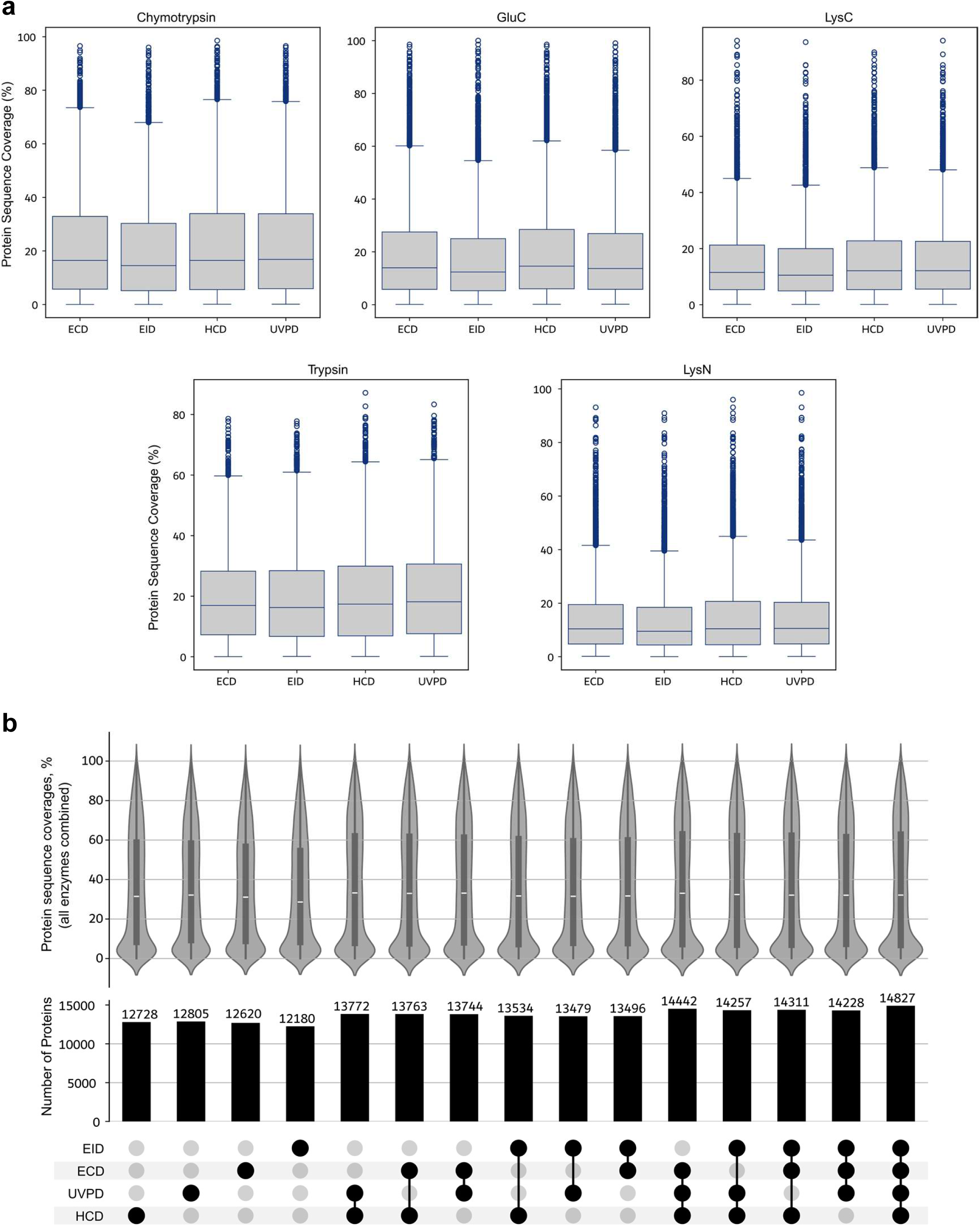
Proteome coverages in DDA data. **a**, Box plots showing the distributions of protein sequence coverages of all proteins identified in different enzyme datasets using different fragmentation techniques. **b,** Upset plots showing the distributions of protein sequence coverages of all proteins in all enzymes combined in various combinations of fragmentation methods.

